# The rise and fall of the photoinhibition-related energy dissipation q_I_

**DOI:** 10.1101/2021.03.10.434601

**Authors:** Wojciech J. Nawrocki, Xin Liu, Bailey Raber, Chen Hu, Catherine de Vitry, Doran I. G. Bennett, Roberta Croce

## Abstract

Photosynthesis converts sunlight into chemical energy, sustaining the vast majority of the biosphere. Photosystem II (PSII), the oxygen-forming enzyme that initiates photosynthesis, is however particularly prone to light-induced damage in a process known as photoinhibition, which limits the productivity of both aquatic and land photosynthesis. Photoinhibition is associated with an energy dissipation process of unknown origin, termed q_I_. Here, we present a detailed biophysical and biochemical in vivo study of q_I_ in model green alga Chlamydomonas reinhardtii. Time-resolved fluorescence measurements demonstrate the origin of q_I_, and indicate the PSII reaction centre as the site of the quencher. Oxygen-dependence of quenching site formation, but not photoinhibition itself, is shown, suggesting that two types of PSII damage – donor and acceptor-side impairment – can be separated. We then demonstrate that the quenching loss takes place in the absence of PSII repair, and is mediated by the degradation of photoinhibited PSII cores by the FtsH protease. Finally, we integrate data ranging from picoseconds to hours in the context of structure-function excitation energy-transferring membrane patches, revealing the extent of PSII heterogeneity from the onset of photoinhibition until the breakdown of damaged PSII.

**Graphical Abstract:** 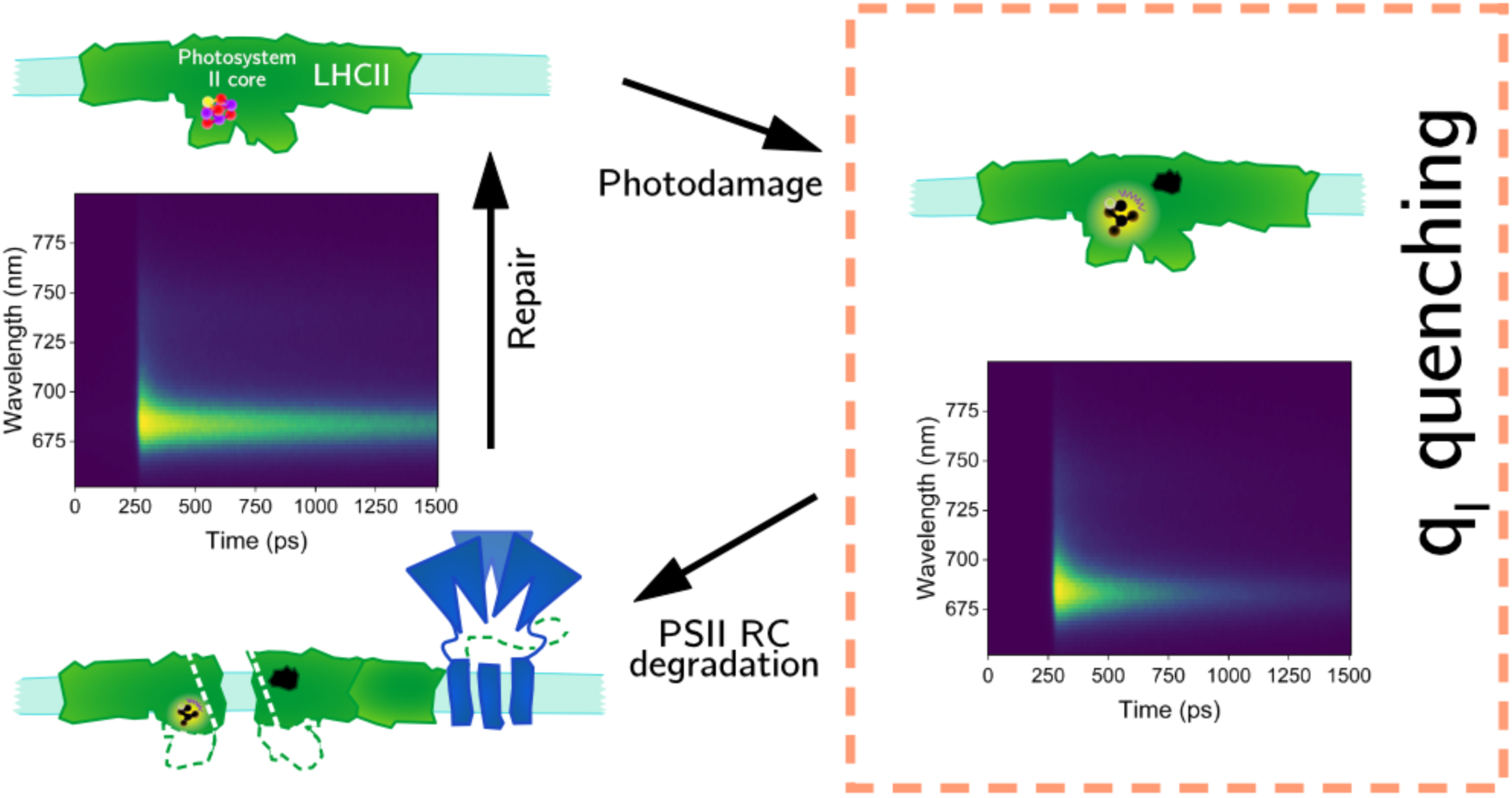

**Highlights:**

- Upon photoinhibition, oxygen sensitization results in an irreversible formation of quenching (q_I_) and inactivation of Photosystem II
- q_I_ takes place in the PSII reaction centre
- Photoinhibition-induced D1 cleavage is much slower than q_I_ formation
- FtsH metalloprotease is required to degrade quenching PSII reaction centres
- A multiscale energy transfer model describes heterogeneity of PSII during photoinhibition

## Introduction

Oxygenic photosynthesis supplies virtually the entire biosphere with chemical energy and oxygen. The first steps of this process involve light harvesting and H_2_O oxidation by Photosystem II (PSII), the water:plastoquinone photooxidoreductase. PSII is arranged in supercomplexes in the appressed regions of the thylakoid membranes within plant and algal chloroplasts (Daum et al., 2010; Wietrzynski et al., 2020). In the model green alga *Chlamydomonas reinhardtii* these multimeric, pigment-protein supercomplexes consist of i) PSII core: composed of the reaction centre (RC) complex (the D1 and D2 proteins, cytochrome *b*_559_ and PsbI), the CP43 and CP47 antennae, the luminal proteins PsbO, P, and Q, stabilising the oxygen-evolving Mn_4_CaO_5_ cluster (OEC), and 10 other small transmembrane subunits; ii) the minor antennae CP26 and CP29; and iii) the light-harvesting complex II (LHCII) (Croce and van Amerongen, 2020; Shen et al., 2019). The antennae greatly enhance the absorption cross-section of PSII allowing efficient light harvesting and excitation energy transfer to the reaction centre, where charge separation takes place and electron transfer begins. The antenna complexes also link different RCs, allowing the excitation to travel in the membrane until it is used to drive stable photochemistry (Bennett et al., 2018; Joliot and Joliot, 1964). However, both the charge-separated states and the long-lived chlorophyll excitation are potentially detrimental to PSII due to oxygen sensitization. PSII has adapted to minimise reactive oxygen species (ROS) formation thanks to the heterodimeric RC complex design (Brinkert et al., 2016; Johnson et al., 1995; Rutherford et al., 2012) and through auxiliary mechanisms that dissipate a fraction of the excitation (Peers et al., 2009; Roach et al., 2020). The latter principally describes energy quenching, termed q_E_ NPQ (Non-Photochemical Quenching), which in Chlamydomonas is mediated by the LHCSR3 antenna after protonation of its lumen-exposed residues (Bonente et al., 2011; Liguori et al., 2013; Peers et al., 2009; Tian et al., 2019). Additionally, PSII regulates its absorption cross-section via the process of State transitions, a phosphorylation-dependent antenna redistribution between PSII and PSI (Bonaventura and Myers, 1969; Croce, 2020; Murata, 1969; Nawrocki et al., 2016). These mechanisms can decrease the probability of PSII damage, known as photoinhibition.

Known for more than six decades, photoinhibition is a process of slowly-recovering, light-induced decrease of PSII activity (Kok, 1956). It causes inactivation of the RC and damage to the D1 protein (Kale et al., 2017; Ohad et al., 1984), which is then followed by degradation and repair with de novo synthesis of PSII subunits (Aro et al., 1993; Hippler et al., 1998; Murata et al., 2007; Nixon et al., 2010, 2010; Tyystjärvi, 2013; Vass and Cser, 2009). Crucially, photoinhibition is a major limiting factor to both terrestrial and aquatic photosynthesis (Chen et al., 2020; Long et al., 1994), and PSII function impairment is particularly strong when high light is combined with other environmental stresses (Murata et al., 2007). It is thus expected to increase as a result of climate change and negatively impact crop productivity (Ainsworth and Ort, 2010).

As demonstrated in the seminal works of Tyystjärvi and Murata groups, a two-step process explains PSII inactivation (Hakala et al., 2005; Ohnishi et al., 2005; Tyystjärvi, 2013). The donor side of PSII (the OEC) becomes damaged independently of PSII photochemistry, when cells are exposed to blue and UV light. On the other hand, acceptor side damage depends on the rate of photosynthetic reactions, and can be partly alleviated by photoprotection mechanisms. Finally, in both cases, NPQ allows PSII repair to proceed efficiently thanks to a decrease of ROS formation (Hakala et al., 2005; Ohnishi et al., 2005; Roach et al., 2020). Photoinhibition thus illustrates the dilemma between optimisation of photosynthesis and excess energy management.

Surprisingly, photoinhibition itself is associated with a decrease in chlorophyll fluorescence yield (Krieger et al., 1992; Matsubara and Chow, 2004; Richter et al., 1999; Zavafer et al., 2019). This behaviour is unexpected because in other conditions where PSII is inactive or absent (e.g. in the absence of electron acceptors (Baker, 2008; Kautsky and Hirsch, 1931); in the presence of the herbicide DCMU blocking PSII acceptor side (Lazár, 1999); in the PSII RC knock-out mutant (Wollman et al., 1980); and when only LHCs accumulate in the thylakoids (Dinc et al., 2016)), the fluorescence level is high (average Chl* lifetime of >1.2 ns, e.g. (Tian et al., 2019)), due to the lack of photochemical quenching from charge-separating PSII. The site of q_I_, the quenching species, the mechanism, and the role of the energy dissipation related to photoinhibition, are at present unknown.

To study photoinhibition and the related quenching, we used an integrated approach *in vivo*. Multiscale analysis of fluorescence changes from picoseconds to hours provided information about q_I_ and the extent of photodamage of the photosynthetic apparatus. The use of a range of mutants allowed us to pinpoint the location of q_I_ and the mechanism of its formation and loss. Finally, we used membrane-scale modelling of energy excitation transfer to reveal the extent of heterogeneity in PSII populations upon photoinhibition.

## Results

### Loss of fluorescence under high light treatment

The protocol used throughout the study to investigate the origin of the slowly-reversible chlorophyll fluorescence decrease upon high light (HL) treatment in Chlamydomonas is shown in Fig. 1A. To focus exclusively on the photoinhibition-related effect, the experiments were performed in the absence of q_E_ (using cells not previously exposed to HL), PSII repair (in the presence of lincomycin), and initially in the *stt7-9* strain, which is unable to perform State transitions (Depège et al., 2003).

**Figure 1.**
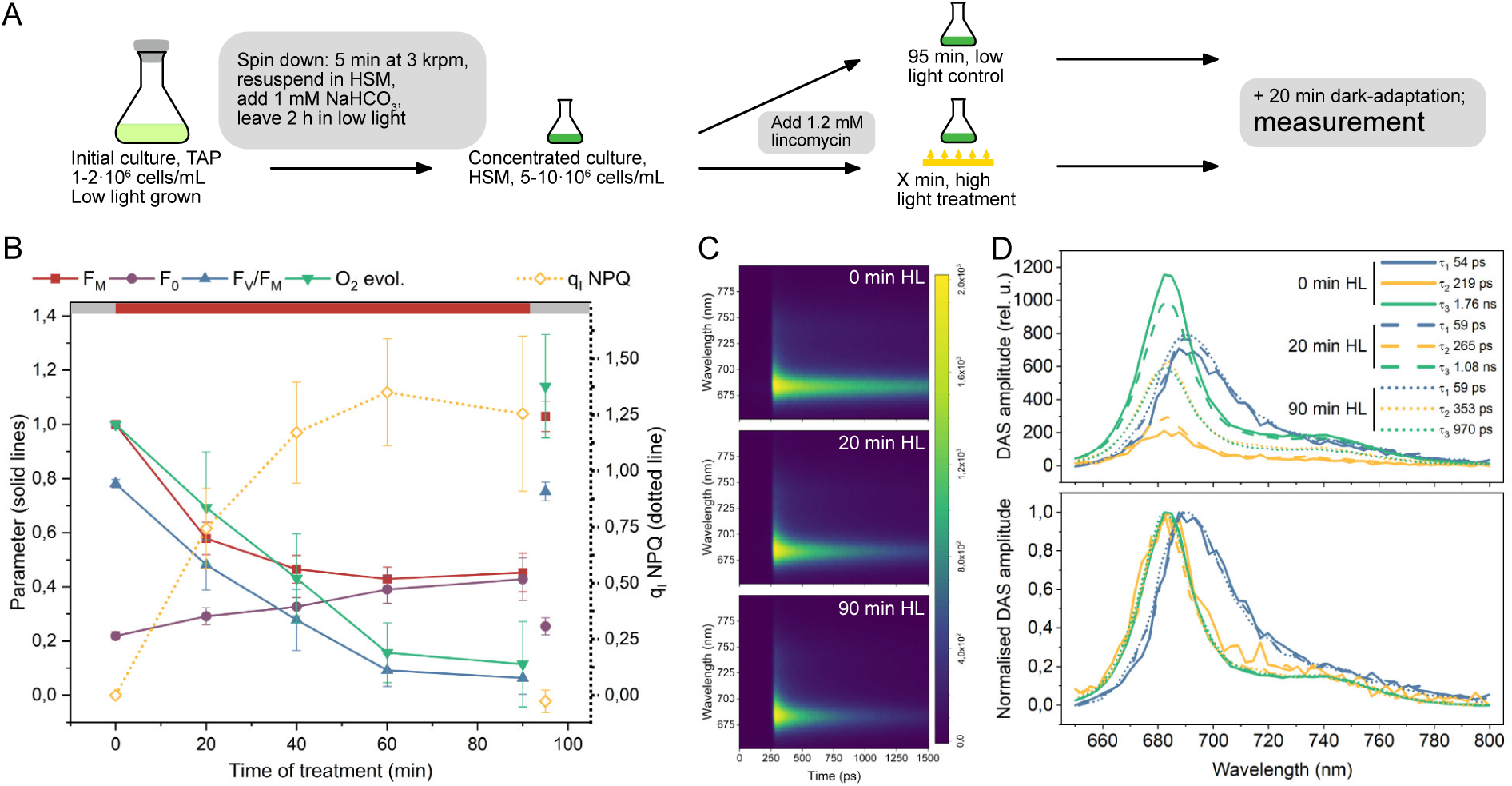
Decrease of chlorophyll fluorescence following photoinhibition in vivo is a quenching. (A) scheme of the treatment prior to the measurements. (B) quantification of the changes in relevant parameters throughout the HL treatment in the *stt7-9* strain. The last timepoint (95 mins) represents the value in samples treated with lincomycin but not exposed to HL. n = 6 ± S.D. (C) Two-dimensional maps of time-resolved fluorescence data of the *stt7-9* strain at t = 0, 20 and 90 min of HL treatment, detected with a Streak camera setup. False colouring depicts the intensity (number of photons) in each bin. A representative dataset is shown. (D) Decay-associated spectra (DAS) of the data in (C). Top panel, the sums of spectra integrals at t = 20 and 90 min were normalised to the t = 0 sum. Lifetimes and spectra were free parameters of linked fitting of 3 biological replicas of the experiment (in total 9 images; see analysis of the other replicas in Fig. S15 and fitting quality in Figs S16-S18).

Throughout the 90 minutes of HL treatment (1500 μmol photons / m^2^ / s), the maximal fluorescence yield of the cells (F_M,_ where the PSII RCs are in closed state) decreased to below 50% of its initial value (Fig. 1B, S1), while the dark-adapted fluorescence value (F_0_, PSII RCs in open state) increased (Fig. 1B). The latter observation indicates that quenching coincides with the closing of PSII RCs upon photodamage since if only quenching occurred, F_0_ would also decrease. The F_V_/F_M_ parameter – often used to quantify the extent of PSII damage – strongly correlates with the decrease in oxygen evolution, confirming photoinhibition of PSII during the HL treatment. Photosystem I (PSI) was instead little affected by the treatment, as demonstrated by the small decrease of the amplitude of photooxidisable P_700_ (Fig. S2). Fluorescence quenching is a term that describes an increase in the overall rate of non-radiative excited state decay. It needs to be distinguished from a decrease of fluorescence caused by a decrease in absorption. The observed reduction in F_M_ is on a timescale of < 2 h an energy quenching process, as indicated by the decrease in fluorescence lifetime of the cells (Figs 1C, S3), and the fact that the absorption capacity of the cells decreases only slightly during the HL treatment (Fig. S1). This quenching process is thus hereafter termed q_I_.

### Site of the q_I_ quenching

There exist several possible quenching sites in the thylakoid membranes of Chlamydomonas. These include PSI (which can act as a quencher of PSII when the two complexes are in close contact (Bag et al., 2020); (2) LHCII (as proposed for q_E_ in vascular plants (Horton et al., 2005); (3) LHCSR (the site of the pH-dependent quenching (Tian et al., 2019); and (4) the PSII core itself (through an unknown mechanism). To identify the q_I_ site, a combination of genetic and spectroscopic approaches was employed. First, time-resolved fluorescence measurements were performed before- (0 min HL) and after 20- and 90 minutes of HL treatment to investigate photoinhibition-dependent changes in spectra and lifetimes of the cells (Fig. 1C). Three decay components were sufficient to describe the fluorescence kinetics at each timepoint. The Decay Associated Spectra (DAS) are shown in Fig 1D. The two longer components, with lifetimes τ_3_ = 1.76 ns and τ_2_ = 219 ps before photoinhibition are associated with PSII, while the shortest component had spectrum and lifetime (τ_1_ = 54 ps) typical of PSI. After 90 min HL treatment, the amplitude and lifetime of the longer PSII component decreased, and those of the shorter PSII component increased. Crucially, no noticeable differences were observed in their spectra (Fig. 1D). The absence of new emitting species after HL treatment, in particular one with a red-shifted spectrum, suggests that LHC aggregation is not the mechanism behind q_I_ (Miloslavina et al., 2008). The PSI spectra and lifetimes were similar before and after HL treatment, indicating that energy spillover from PSII to PSI did not take place during photoinhibition. This conclusion is supported by the similarity of q_I_ amplitude and kinetics in the *stt7-9* and ΔPSI strains (Fig. S4).

To verify if q_I_ involves the LHCs via an aggregation-independent mechanism, we examined three strains with reduced antenna content (Fig. S5). In all mutants, the relation between q_I_ and photodamage was similar to reference strains (Fig. S5), supporting the conclusion that the LHCs are not the site of q_I_. We observed that despite the initial absence of LHCSR3 in the cells, expression of this q_E_ -inducing protein took place during HL treatment (Fig. S5). To verify whether it could contribute to q_I_, we analysed the *stt7-9 npq4* double mutant, where LHCSR3 is knocked out (Peers et al., 2009). In this mutant, the q_I_ amplitude and kinetics remained comparable to the control, excluding the option of LHCSR3-dependent q_I_ (Fig. S5).

The results described above suggest that q_I_ occurs within the PSII core. This is supported by the analysis of the ΔPSII mutant, which shows far less quenching and slower induction kinetics than the control and the ΔPSI mutant (Fig S4).

### Loss of q_I_ is light-independent and does not require chloroplast translation

Next, we investigated the dependency of the quenching from PSII repair by measuring the stability of q_I_ in the long timescale in the absence of PSII repair. We also extended the study beyond the *stt7-9* mutant and included three WT strains with distant genetic backgrounds (Gallaher et al., 2015): CC-124, CC-1009, and CC-1690. The influence of State transitions on fluorescence signals during photoinhibitory treatment was accounted for (See Fig. S3 for details).

Following photoinhibition, all tested strains developed q_I_. While the maximal amplitude varied between 1 and 1.4, in all strains q_I_ reached a maximum and then strongly decreased (Fig. 2A; see Fig. S3E for the *stt7-9* strain data). Surprisingly, this decrease proceeded in the presence of lincomycin, indicating that it was not related to *de novo* PSII synthesis (Fig. 2B) or to a recovery of the PSII function, as also indicated by the fact that F_V_/F_M_ did not change during the fluorescence recovery period (Fig. 2C). Furthermore, the loss of quenching occurred already during the HL period. Neither the duration of the HL treatment (after reaching the peak of q_I_ amplitude), nor whether it was followed by a dark period or low-light treatment, had a significant influence on the loss of q_I_ (Fig. S6). Together, these results indicate that a slow, light-independent process which does not require active chloroplast translation governs the q_I_ relaxation.

**Figure 2.**
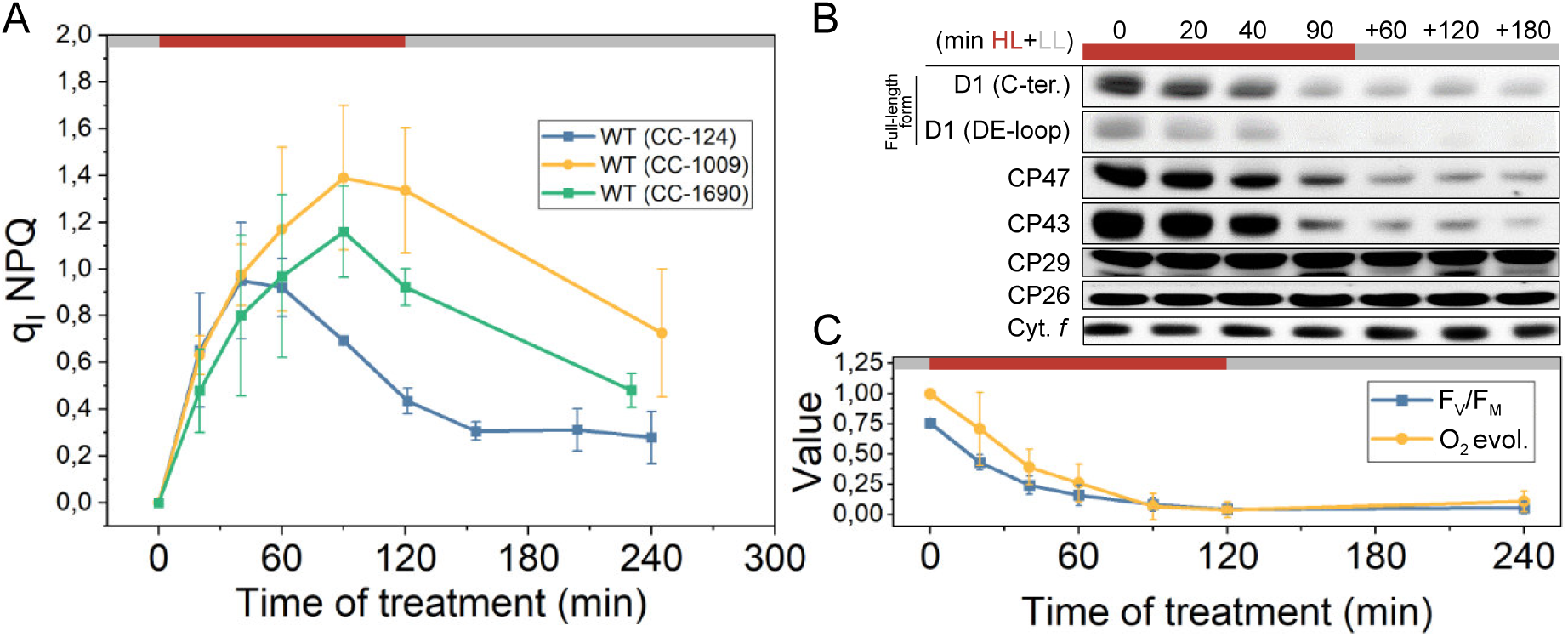
q_I_ is transient and its loss is independent of PSII repair. All experiments shown were performed in the presence of lincomycin, which inhibits chloroplast translation. Red boxes – HL illumination (1500 μmol photons / m^2^ / s); grey boxes – low light period (15 μmol photons / m^2^ / s). (A) Development of q_I_ in three WT strains. (CC-1009, CC-1690: n = 3; CC-124: n = 6) upon exposure of the cells to HL followed by LL. (B) Immunoblotting with antibodies against selected PSII subunits during HL and LL exposure in the CC-124 WT strain. (C) PSII activity in WT (CC-124) during HL treatment followed by a LL period, measured using fluorescence (F_V_/F_M_) and O_2_ evolution capacity. n = 3.

### Loss of q_I_ relies on PSII proteolysis by FtsH

We hypothesized that the slow, light-independent q_I_ loss is due to the degradation of PSII core subunits within which q_I_ occurs (Fig. 2B). To test that, we measured q_I_ in *ftsh1* mutants, where the major metalloprotease involved in PSII degradation (Kato et al., 2012; Malnoë et al., 2014; Wang et al., 2017) is inactive (*ftsh1-1*) or present in a low amount (*ftsh1-3*). The amplitude and kinetics of q_I_ induction in the *ftsh1* mutants closely followed those of the reference WT strain (Fig. 3A, S7). However, the mutants remained quenched for a longer time (Fig. 3A, S7) than the controls. This behaviour was *ftsh1*-dependent, as demonstrated by the fact that the complemented strains showed q_I_ loss (Fig. 3A, S7).

**Figure 3.**
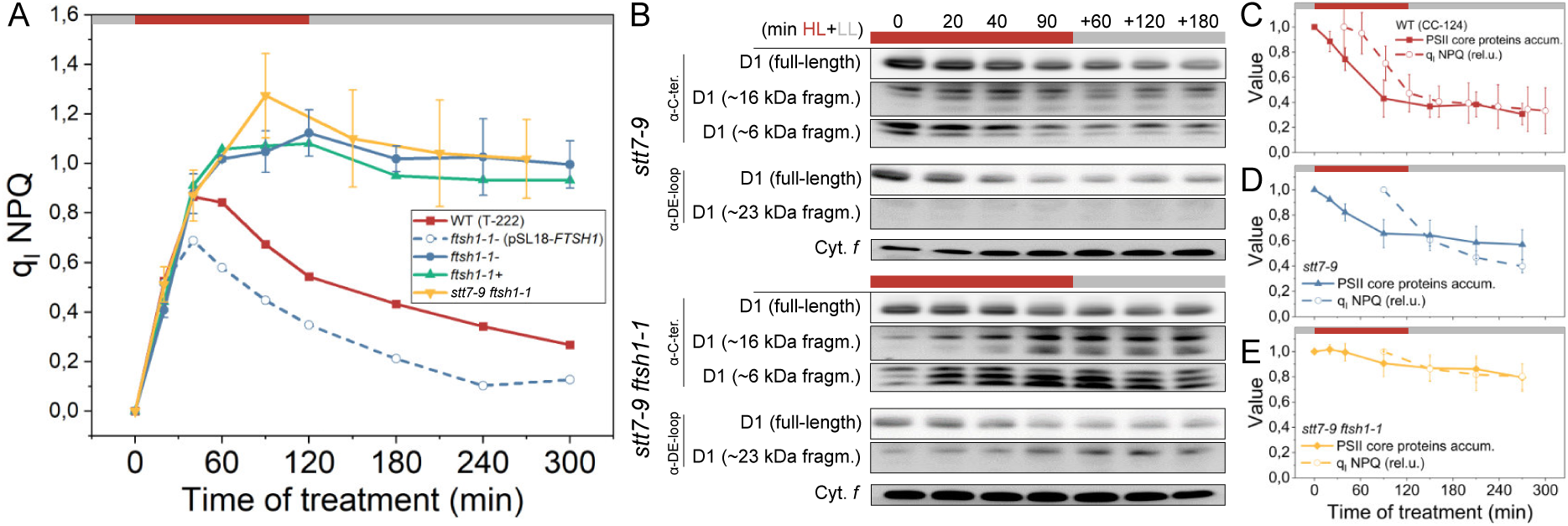
q_I_ quenching loss is due to PSII core proteolysis by FtsH. All experiments shown were performed in the presence of lincomycin, which inhibits chloroplast translation. Red boxes – HL illumination (1500 μmol photons / m^2^ / s); grey boxes – low light period (15 μmol photons / m^2^ / s). (A) q_I_ quenching behaviour upon photoinhibition. WT (T-222) is the parental strain of the *ftsh1-1* mutant. Two separate clones of the *ftsh1-1* allele are shown together with the *ftsh1-1*:pSL1 FTSH1 complemented strain and the double mutant *stt7-9 ftsh1-1*. See also fig. S7 for the *ftsh1-3* allele. (B) D1 degradation timecourse in *stt7-9* and *stt7-9 ftsh1-1* strains, followed by immunoblotting. One representative biological replica is shown; cytochrome *f* accumulation is not affected by photoinhibition and was used as a loading control. (C) PSII RC proteins loss (average of relative signal from 3 antibodies [α-DE loop and α-C-ter. of D1; α-CP47], see Fig. S9 for raw quantification data), n = 1, and relative q_I_ NPQ, n = 6, in WT (CC-124). (D) PSII RC proteins loss, n = 3, and relative q_I_ NPQ, n = 3, in *stt7-9* strain. (E) PSII RC proteins loss, n = 3, and relative q_I_ NPQ, n = 3, *stt7-9 ftsh1-1* strain.

To be able to correlate the protein degradation with fluorescence changes in the absence of State transitions influence on the emission, we constructed the *stt7-9 ftsh1-1* double mutant (Fig. S8). As shown in Fig. 3B, both *stt7-9* and *stt7-9 ftsh1-1* strains exhibited a loss of full-length D1 protein upon HL treatment, with the initial cleavage being slower in the double mutant (Fig. S9). As observed before, only the *stt7-9 ftsh1-1* strain accumulated short fragments of D1 (Fig. 3B), due to the absence of *ftsh*-mediated exoproteolytic degradation (Malnoë et al., 2014). In line with these observations, Blue-native (BN) PAGE showed that the smaller PSII supercomplexes decreased far more in *stt7-9* than in the double mutant during photoinhibition (Fig. S10).

Together, these experiments demonstrate the impaired PSII RC complex proteolysis in the *ftsh1* background. Overall, the loss of q_I_ showed correlation with the PSII proteins degradation across *stt9* and *stt7-9 ftsh1-1* strains (as well as in the WT), although it exhibited a lag with regards to proteolysis (Fig. 3C-E). These observations indicate that loss of intact PSII core leads to the loss of quenching (Fig. 3C-E). Interestingly, the lag in observed quenching loss could potentially be explained in the context of energetic connectivity in the thylakoids as decrease of quenching due to PSII degradation being temporarily limited by presence of quenching side in nearby PSIIs (See section *Photoinhibited PSII reaction centres are heterogenous* for details).

The location of the q_I_ site in the photoinhibited PSII core was finally confirmed by the ΔPSII mutant, in which no loss of q_I_ was observed neither in HL nor in LL, indicating that the slow quenching observed in this strain (Fig. S4) has a different origin, and resembled what was before observed in lacking both photosystems cells exposed to HL (Tian et al., 2015).

### Formation of q_I_ site requires oxygen

To understand the sequence of events upon photoinhibition we focused on the initial kinetics of q_I_ and other functional- and biochemical obervables. As shown in Fig. 4A, loss of fluorescence (F_M_) was by far the fastest event upon HL treatment (t_1/2_ of ∼15 min). F_V_/F_M_ decreased slower (t_1/2_ of ∼30 min), but as expected it showed good correlation with the decrease of oxygen evolution. Crucially, the degradation of PSII RC proteins was significantly slower than the changes in fluorescence and oxygen evolution (Fig. 4A), in line with previous reports (Aro et al., 1993). These results substantiate the above hypothesis that protein degradation is related to the loss of q_I_ rather than its induction, and highlight that q_I_ is one of the earliest events upon photoinhibition.

**Figure 4.**
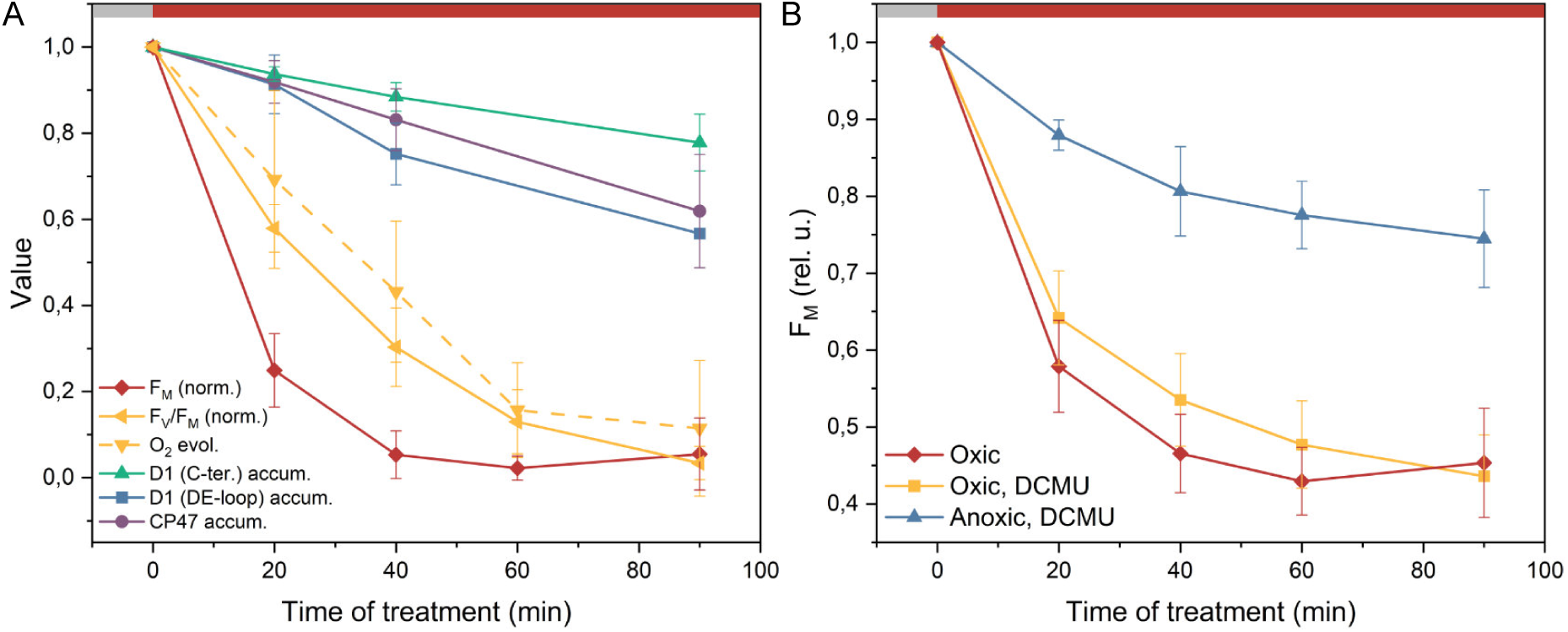
q_I_ site formation is rapid and requires the presence of oxygen. All experiments were performed in the presence of lincomycin, which inhibits chloroplast translation, in the *stt7-9* strain. Red boxes – HL illumination (1500 μmol photons / m^2^ / s); grey boxes – low light period (15 μmol photons / m^2^ / s). (A) Changes in photosynthetic parameters and protein accumulation upon photoinhibition. F_M_, F_V_/F_M_, and oxygen evolution changes n = 11; PSII RC proteins accumulation, n = 3. F_M_ was normalised between its maximal and minimal value, and F_V_/F_M_ to its maximum. (B) Fluorescence quenching in anoxic conditions. n = 3 (DCMU; anoxic, DCMU) and n = 11 (oxic conditions).

The fact that the formation of the quenching site within the core of PSII precedes D1 cleavage leaves several options regarding the quenching mechanism, including non-photochemical energy dissipation and charge separation-based quenching. It was recently shown that protein oxidation events take place early upon photoinhibition (Kale et al., 2017) and that they can induce quenching in pigment-protein complexes (Lingvay et al., 2020); furthermore, it was observed that oxygen influences the F_M_ level during photoinhibition (Gong and Ohad, 1991). We thus investigated whether oxygen is necessary for (1) the quenching itself, or (2) for the formation of the q_I_ site. To test the first hypothesis, we induced anoxia in photoinhibited cells. In this case, the quenching capacity remained unchanged, demonstrating that O_2_ is not necessary for energy dissipation (Fig. S11). To test the second hypothesis, photoinhibitory treatment was conducted in anoxia. Addition of DCMU was necessary to prevent O_2_ evolution by PSII but it did not affect the loss of fluorescence (Fig. 4B). The q_I_ amplitude was strongly reduced in anoxic conditions, (Fig. 4B) and the PSII closure was slower (Fig. S12), suggesting that ROS-mediated oxidation of a specific pigment within the PSII RC creates the quenching site (Fig. 4B). In conclusion, the oxygen-dependence of q_I_ site formation indicates that quenching arises by the inactivation of PSII RC, likely through an RC chlorophyll oxidation. These observations support an acceptor side photoinhibition-related formation of the q_I_ site.

### Photoinhibited PSII reaction centres are heterogenous

The induction and loss of photoinhibition-related quenching is a complex process that occurs on minutes to hours timescale. During high light exposure, changes in fluorescence parameters, such as F_M_ and F_V_/F_M_, occur faster than the degradation of the reaction centre proteins (Fig. 2-4). Notably, the decrease of F_M_ precedes the loss of PSII activity (F_V_/F_M_). Moreover, during the dark period following high light exposure, there is a loss of q_I_ correlating with the D1 preoteolysis after a lag phase and without a corresponding change in F_V_/F_M_ (Fig. 2, 3 C-E). To understand these observations in the context of a structure-based model of light harvesting (Fig. 5 A-C), we hypothesized that there are two distinct types of photoinhibited RCs, both unable to perform photochemistry: (1) ‘quenching’ centres which exhibit q_I_ capacity and (2) ‘broken’ centres which do not. All together our minimal model contains active RCs that are either ‘open’ (F_0_) or ‘closed’ (F_M_), and inactive RCs that are either ‘quenching’ or ‘broken’ (Fig. 5D).

**Fig. 5.**
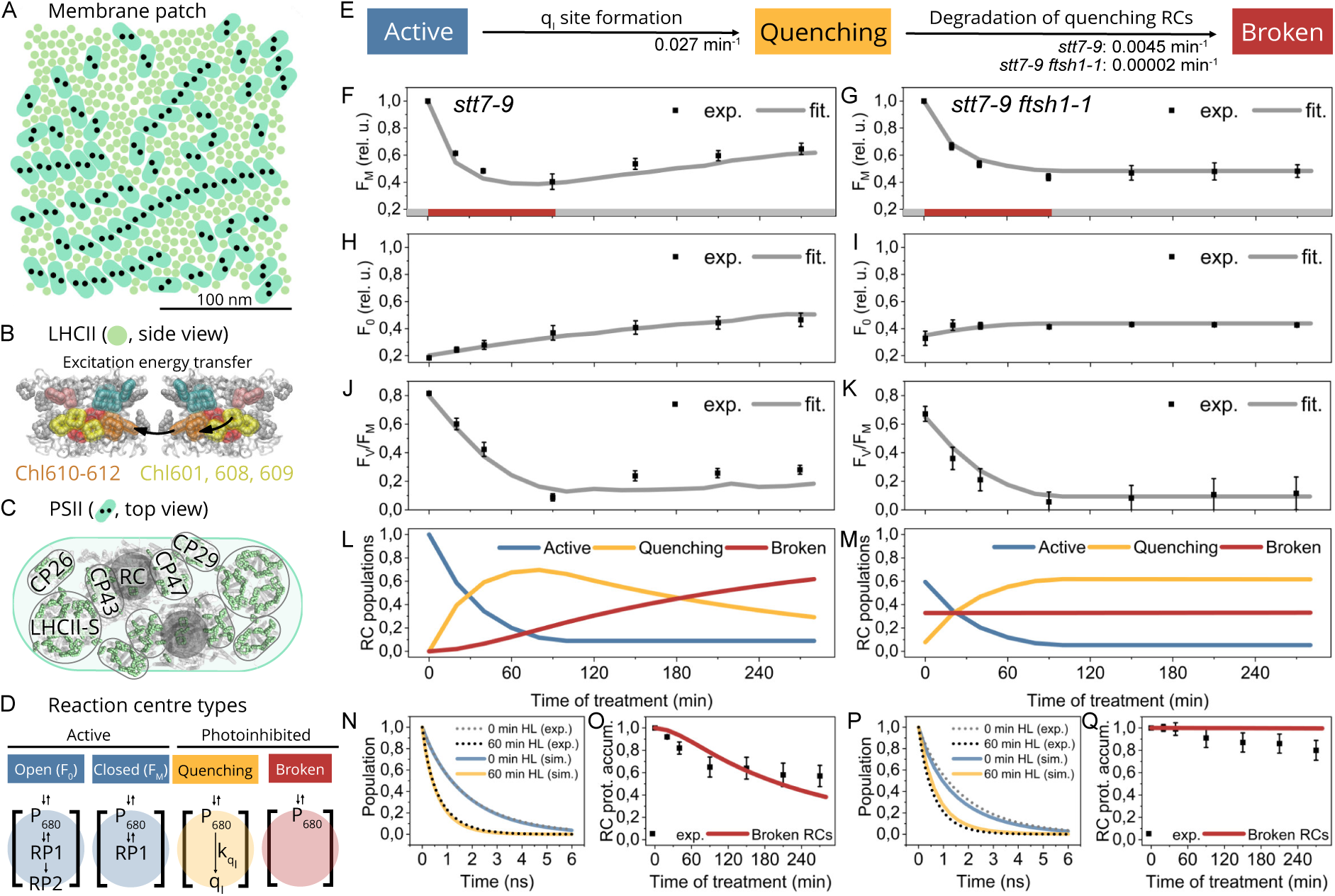
Heterogeneity of photoinhibition as revealed through membrane-scale structure-function modelling. The model represents an appressed region of the thylakoid membrane containing PSII-LHCII supercomplexes. In all steady-state data, the constant ∼3% fluorescence contribution from PSI was subtracted, and the single PSI component (∼60 ps) was omitted from in time-resolved fluorescence plots. Red boxes – HL illumination (1500 μmol photons / m^2^ / s); grey boxes – low light period (15 μmol photons / m^2^ / s). (A) Organisation of photosynthetic complexes in the appressed region-like membrane patch used for the modelling. (B) Coarse-graining of pigment domains in LHCII. Foerster resonance energy transfer between the domains within an LHCII and between adjacent antenna is represented with arrows. (C) PSII supercomplex structure used in the model (Caffarri et al., 2009). (D) Types of RCs used in modelling and their characteristics. An ‘open RC’ has both an initial reversible charge separation (RP1) and a slower irreversible charge separation (RP2). A ‘closed RC’ has a slower rate of initial charge separation (RP1) with a faster rate of recombination compared to open RCs and does not have the ability to perform irreversible charge separation. A ‘quenching RC’ has no rate of charge separation but has a separate non-photochemical quenching mechanism with a rate k_qI_. A ‘broken RC’ has no charge separation or quenching pathways. (E) Kinetic model of photoinhibition used for the modelling. The rates were obtained by fitting the data from panels F-K. (F, G) Experimental (points (mean) ± SD; n = 3) and modelled fluorescence yields (thick line) during photoinhibition when all RCs are closed (F_M_ state) in *stt7-9* (F) and *stt7-9 ftsh1-1* strains (G). (H, I) Experimental (points (mean) ± SD; n = 3) and modelled fluorescence yields (thick line) during photoinhibition when all active RCs are open (F_0_ state) in *stt7-9* (H) and *stt7-9 ftsh1-1* strains (I). (J, K) Experimental (points (mean) ± SD; n = 3) and modelled changes in the F_V_/F_M_ parameter (thick line) during photoinhibition in *stt7-9* (J) and *stt7-9 ftsh1-1* strains (K). Note that the small recovery of F_V_/F_M_ is due to a shorter time of photoinhibition used compared to Fig. 3. (L, M) Modelled kinetics of the populations of active-, quenching-, and broken RCs during photoinhibitory treatment in *stt7-9* (L) and *stt7-9 ftsh1-1* strains (M) (N, P) Experimental (dotted line; representative repeat shown) and modelled (continuous line) time-resolved fluorescence traces of PSII-related components in *stt7-9* (N) and *stt7-9 ftsh1-1* strains (P). (O, Q) Experimental (points (mean) ± SD; n = 3) and modelled (thick line) changes in the accumulation of full-length RC proteins (D1 and CP47) throughout photoinhibition in *stt7-9* (O) and *stt7-9 ftsh1-1* strains (Q).

We built a kinetic model to describe the slow (min-hour) processes that convert reaction centres between the four types (Fig. 5E). We explain the decrease of F_M_ following high light exposure by assuming there is a light-dependent reaction that converts active RCs to quenching RCs. We assume the rate of q_I_ site formation is equivalent in both the *stt7-9* and the double mutant *stt7-9 ftsh1-1* (see supplementary discussion for other models). In the absence of PSII repair, the loss of fluorescence quenching suggested a proteolytic degradation of quenching RC complexes (Fig. 3), which we propose forms ‘broken’ RCs. The *ftsh1* mutation decreases the proteolytic activity and thereby reduces the conversion rate of quenching into broken complexes. Thus, our kinetic model has three parameters to describe the time evolution of reaction centre populations: the rate from active to quenching (‘q_I_ site formation’) and the rate from quenching to broken (‘degradation of quenching centres’), which has two possible values depending on the presence/absence of FtsH.

We connected the evolving reaction centre populations to the fluorescence observables using a structure-based model of excitation-energy transport on a 40000 nm^2^ patch of the appressed thylakoid membrane (Amarnath et al., 2016; Bennett et al., 2013, 2018) (Fig. 5A). Excitation-energy transfer between chlorophyll domains was described using generalized Forster theory (Fig. 5B, C; see methods for details), (Scholes, 2003; Sumi, 1999) and charge separation at active reaction centres ((Bennett et al., 2013; Caffarri et al., 2009); Fig. 5C) was described using previously established parameters (see Methods and (Amarnath et al., 2016; Bennett et al., 2013) for details). The rate of excitation dissipation (k_qI_) at quenching RCs was treated as a free parameter due to the absence of any experimental bounds. For a given population of open, closed, quenching, and broken centres we constructed the ensemble average fluorescence values using twenty realizations of different random assignments of PSII complexes to each group. The simulated ensemble average was then compared to the experimental fluorescence measurements.

We constructed the best fit model for photoinhibition by simultaneously minimizing the error compared to the steady-state fluorescence measurements for both the *stt7-9* and *stt7-9 ftsh1-1* mutants (Fig. 5E). For *stt7-9* at 0 min HL, the excitation-energy transfer model successfully reproduces both steady-state F_V_/F_M_ (sim.: 0.8, exp.: 0.82) and time-resolved fluorescence measurements (Fig. 5N). The *stt7-9 ftsh1-1* double mutant at 0 min HL has a substantially shorter F_M_ fluorescence lifetime compared to the *stt7-9* strain, which we assume arises from previous exposure to the growth light and slow PSII repair (Kato et al., 2012; Malnoë et al., 2014). We fit the short F_M_ fluorescence lifetime to a residual mixture of broken (34%) and quenching (6%) RCs (Fig. 5P), which also reproduces the steady-state decrease in F_V_/F_M_ value (sim.: 0.65, exp.: 0.67) at 0 min HL.

The best fit kinetic model has a q_I_ quenching rate (k_qI_) of 0.1 ps^-1^. A single rate of q_I_ site formation (0.027 min^-1^) successfully describes the onset of quenching in both strains, and shows a rate of quenching centres degradation that is 2 orders of magnitude times slower in *stt7-9 ftsh1-1* compared to *stt7-9* (Fig. 5E-M). The model further supports the simultaneous irreversible closing of RCs and q_I_ site formation. Overall, the photoinhibition model proposed here not only correctly simulates the steady-state fluorescence in both strains with a minimal kinetic model, but also reproduces the trends of protein degradation (Fig. 5O, Q). The latter thus suggests that the proteolytic degradation of RC complexes is sufficient and necessary to abolish q_I_, and that the lag observed in Fig. 3D and E could indeed be simply interpreted as a result of energetic connectivity. We finally confirmed the self-consistency of this model by reproducing fluorescence lifetime measurements after 60 min HL treatment (Fig. 5N, P).

Here we have demonstrated that a minimal model containing two types of photoinhibited RCs and three kinetic rates can reproduce both fluorescence measurements reporting on the ultrafast process of light harvesting and immunoblotting data describing protein degradation on the minutes timescale (see Supplementary Discussion and Fig. S19 for more details). Therefore, the model links kinetics spanning 14 orders of magnitude and provides a predictive framework for understanding a broad range of photoinhibition-related events in vivo. Crucially, this model is parsimonious, successfully accounting for the q_I_ formation- and RC closure upon photodamage, without a necessity of evoking two quenching sites and mechanisms proposed earlier (Matsubara and Chow, 2004; Zavafer et al., 2019).

## Discussion

It is of little surprise that photoinhibition, damaging one of the most complex enzymes in biology, is itself a complicated process. Understanding PSII damage and repair is nonetheless crucial due to the importance of photoinhibition in crop and aquatic photosynthesis (Ainsworth and Ort, 2010; Chen et al., 2020; Long et al., 1994; Murata et al., 2007). However, the molecular nature of q_I_ has been rarely addressed in the literature (Krieger et al., 1992; Matsubara and Chow, 2004; Richter et al., 1999; Zavafer et al., 2019). In this work we have simultaneously analysed several aspects of this process which will be discussed here to provide a picture of the formation and relaxation of q_I_ (Fig 7).

### Connectivity between PSII complexes and its influence on F_V_/F_M_

Historically, oxygen evolution, spectroscopy, and biochemical measurements were used to quantify PSII damage (Chow and Aro, 2005; Hippler et al., 1998; Tyystjärvi, 2013). In particular, changes in the photochemical yield of PSII assessed through fluorescence measurements (e.g. F_V_/F_M_) are commonly used to quantify photoinhibition. However, the decrease of F_V_/F_M_ is not linearly related to the accumulation of broken PSIIs, but rather it describes the global maximal photochemistry. This is due to the energetic connectivity, which results in an increase of the antenna size of the active centres by pigments belonging to the photoinhibited complexes (up to 50 nm apart (Amarnath et al., 2016)). Our simulations indicate that formation of damaged centres is faster than the decrease of F_V_/F_M_ and slower than the apparent F_M_ changes (Fig. 6), in line with previous experiments (Oguchi et al., 2009).

**Fig. 6.**
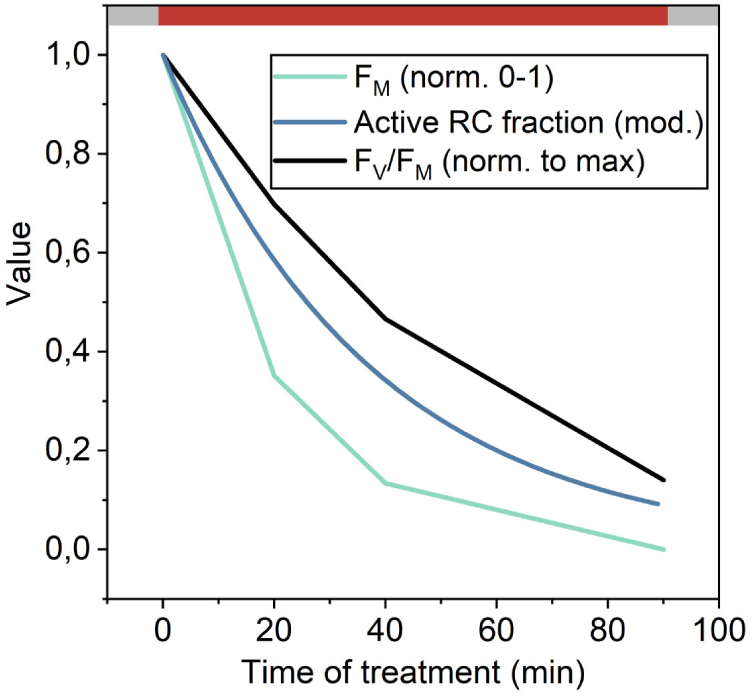
Kinetics of fluorescence changes and the variation in active RC concentration upon photoinhibition. Concentration of the active RCs obtained from membrane-scale simulations. Means of fluorescence measurements in the *stt7-9* strain are depicted, n = 3. F_V_/F_M_ was normalised to its maximal value, F_M_ between 0 and 1. Red box – HL illumination (1500 μmol photons / m^2^ / s); grey boxes – low light period (15 μmol photons / m^2^ / s).

**Figure 7.**
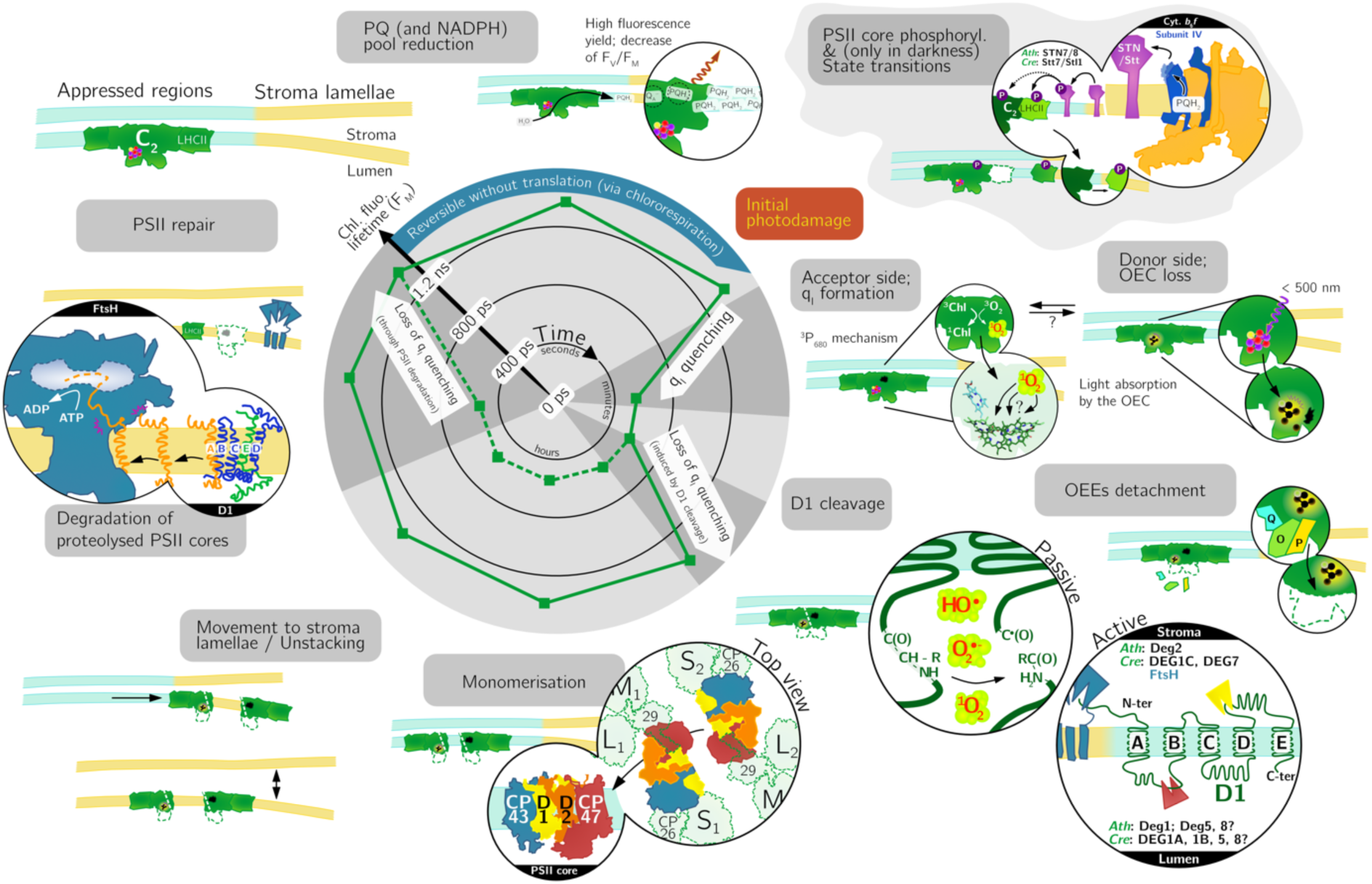
Schematic model of the induction and relaxation of q_I_ upon photoinhibition. Initial photodamage to PSII supercomplex (core dimer, C_2_; and antenna, LHCII) can be described by two mechanisms: the excess-light induced ROS formation and oxidation of PSII subunits and pigments, as well as the direct OEC damage (Tyystjärvi, 2013). q_I_ (inner polar plot of the average fluorescence lifetime in vivo), established with halftime of ∼15 minutes, is concomitant with photodamage. As a result of PSII RC complex degradation the quenching is lost and the lifetime of fluorescence increases back to the initial F_M_ state. Following D1 cleavage the PSII monomerises but remains at least partly assembled despite the endoproteolysis of its reaction centre protein, and translocates to non-appressed regions of the thylakoids (or else the appressed regions unstack) (Barbato et al., 1992; Hundal et al., 1990). Transmembrane thylakoid FtsH protease degrades the PSII RC complex. PSII repair with de novo synthesis of at least D1 subunit takes place to re-establish the initial state.

### Mechanism of q_I_ and potential heterogeneity of photodamage

It is now generally accepted that a two-step model best explains the initial PSII photodamage (Hakala et al., 2005; Ohnishi et al., 2005; Tyystjärvi, 2013). First, direct absorption of blue or ultraviolet light by the manganese ions of the OEC damages the donor side of PSII. Then, donor side-impaired PSII trigger changes on their acceptor side. These changes trigger protein oxidation (Kale et al., 2017) and cleavage (Aro et al., 1993).

Our results agree with the above model, but indicate that the two steps take place in parallel rather than sequentially, in agreement with a previous proposal (Oguchi et al., 2009, 2011). The oxygen dependence of q_I_ formation (Fig. 4) and the slower F_V_/F_M_ decrease in anoxia (Fig. S12) suggest that in our experimental conditions the acceptor-side damage is faster than the donor-side photoinhibition, but that in the absence of O_2_ the latter still takes place. Crucially, photoinhibition linearly correlates with light intensity ((Tyystjärvi, 2013); Fig. S13) and RC closure takes place simultaneously with the q_I_ site formation (Fig. 5). We therefore consider that the acceptor-side damage involves oxidation of PSII RC pigment(s) through in situ singlet oxygen sensitization proceeding by PSII charge recombination. As demonstrated by Rehman et al. (Rehman et al., 2013), ^1^O_2_ formation by ^3^P_680_ (a by-product of charge recombination) linearly correlates with light intensity, in line with this hypothesis (Vass and Cser, 2009). The parallel model implies that the two steps of photoinhibition can be separated. An argument in agreement with that is provided by the anoxic photoinhibition experiment: we observed the appearance of a light-induced quenching of small amplitude (Fig. S11) in samples which undergone HL treatment. This quenching hints at a capacity of P_680_^+^ formation due to the dysfunctional donor side, and thus a partially active acceptor side thanks to the absence of oxygen sensitization. The dependence of photoinhibition and fluorescence parameters on light colour support the same conclusion (Oguchi et al., 2009).

Interestingly, an intrinsic regulation of the rate of ^1^O_2_ production is built into the PSII, decreasing the rate of PSII charge recombination (Johnson et al., 1995; Rehman et al., 2013). It depends both on the presence of functional donor- and acceptor sides of PSII (Brinkert et al., 2016; Johnson et al., 1995). The midpoint redox potential (*E_m_*) of Q_A_ quinone increases by around 115 mV when OEC is impaired and by 75 mV in the absence of the HCO3 ^-^ between Q_A_ and Q_B_. As a consequence, direct recombination between P_680_^+^/Q_A_^-^ (or Tyr_Z_^+^/Q_A_^-^) is favoured over a repopulation of the P_680_* state. Crucially, such charge recombination could in principle also constitute the q_I_ mechanism – when the donor side of PSII is impaired upon photoinhibition, radiationless recombination would result in a quenching of fluorescence (Krieger et al., 1992). Our experiments, however, show that it is an unlikely scenario. For the recombination mechanism to work, an oxidised Q_A_ quinone is necessary, while anoxia induced post-photoinhibition – which non-photochemically reduces this cofactor (Mus et al., 2005) – had no influence on the quenching. Conversely, oxygen presence was necessary to form the q_I_ site within the PSII reaction centre. Formation of ^1^O_2_ is well established upon photoinhibition and ROS have been shown to damage the protein scaffold (Kale et al., 2017). Pigment oxidation also results in a shortening of chlorophyll excited state lifetime (Lingvay et al., 2020). Finally, singlet oxygen concentration increases linearly with light intensity, as does the rate of q_I_ formation (Fig. S13). Thus, a hypothesis where ^1^O_2_ formed via ^3^P_680_ directly oxidises one of the RC pigments, impairing charge separation capacity and forming the q_I_ site, is consistent with all the observations.

The kinetic model of photoinhibition (Fig. 5) provides insights into the heterogeneity of PSII RC composition during HL treatment. Importantly, it demonstrates that even in the case of a null F_V_/F_M_ value, a combination of quenching and broken RCs is present in the thylakoids (Fig. 5L, M). Nonetheless, our model supports a homogenous damage mechanism, with a simple conversion of active RCs to quenchers, followed by their degradation and loss of q_I_ sites (but see also Supplementary discussion and Fig. S19). This finding succeeds in describing notably the kinetically different behaviour between F_M_ and F_V_/F_M_. Additionally, the close match between the appearance of broken centres and PSII core subunits degradation suggests that the latter process abolishes the quenching capacity in vivo.

### Is photoinhibition photoprotective?

Quenching is often associated with photoprotection. In the case of photoinhibition-related quenching, the situation is more complex, as damage is a prerequisite for quenching. The linear dependence of photoinhibition on light intensity and first-order kinetics of PSII function loss (see discussion in (Santabarbara et al., 2002; Tyystjärvi, 2013) and references therein) support the donor-side damage mediated by the direct light absorption by the OEC. This damage cannot be prevented by quenching, but only by photoprotective adaptations such as screens on the leaf surface (Hakala-Yatkin et al., 2010). This interpretation becomes slightly more nuanced in the scope of the two-step photoinhibition model: the donor-side mechanism might be inevitable, but the acceptor-side impairment could potentially be alleviated. However, it is important to stress that F_V_/F_M_ measurements do not distinguish between these two mechanisms (Fig. S12) - using quenching as the observable might be beneficial for suture studies (Fig. 4).

Ultimately, while PSII photodamage studies are invariably done in the absence of PSII repair, in natural conditions limiting ROS production thanks to q_I_ protects the repair machinery (Nishiyama et al., 2006) and alleviates damage in the steady-state (Roach and Krieger-Liszkay, 2019; Roach et al., 2020). As such, q_I_ can positively influence the recovery from photoinhibition, even if not significantly preventing PSII damage (Santabarbara et al., 2001). In particular, PSII assembly provides cues that agree with the need of PSII acceptor-side photoprotection and ROS formation decrease. The abovementioned changes in Q_A_ *E_m_* were recently shown to dampen the singlet oxygen production before PSII became fully assembled (Johnson et al., 1995; Zabret et al., 2020).

Crucially, however, by damaging the PSII RC complex photoinhibition prevents the formation of long-lived quenching in LHCII (Fig. S4). The duration of quenching in HL in the absence of PSII core (Tian et al., 2015) suggests that it is difficult to relax this quenching which can thus be detrimental to photosynthetic efficiency following HL exposure (as suggested in the case of too-slowly-relaxing q_E_ (Kromdijk et al., 2016)). This in turn highlights that locating q_I_ in the replaceable PSII RC is another feat of PSII, whose design and function do not cease to surprise (Brinkert et al., 2016; Zabret et al., 2020).

### Potential mechanisms of D1 damage

Whether offering substantial photoprotection or not, quenching centres are damaged and thus need to be repaired (Baena-González et al., 1999; Chow and Aro, 2005; Hippler et al., 1998; Li et al., 2018), starting with the degradation of the D1 protein. The mechanism of D1 degradation is debated, with at least three known processes involved.

(1) Passive cleavage of D1 was observed upon HL treatment in vitro (Ke, 2001; Shipton and Barber, 1994), likely involving protein radicals formed upon photoinhibition (Davies, 2003; Kale et al., 2017) or even by P_680_^+^. This mechanism yields similar D1 degradation fragments as photoinhibition in vivo, which is presumed to proceed through (2) DEG-mediated endoproteolysis (Andersson and Aro, 2001; Kapri-Pardes et al., 2007; Theis et al., 2019), followed by FtsH-mediated processing of D1 fragments (Malnoë et al., 2014). (3) Direct degradation of D1 by FtsH was also observed (Chow and Aro, 2005), in line with our data (Fig. S9). However, the light-dependent change in the rate of full-length RC proteins degradation (Figs 3, S9) suggests that a combination of these mechanisms takes place in living cells.

The luminal subunits of PSII (PsbO/P/Q; (Shen et al., 2019)) shield the D1 peptide from proteolysis by DEG. Their dissociation was indeed shown to correlate with D1 cleavage (Eisenberg-Domovich et al., 1995; Hundal et al., 1990), and it could constitute the rate-limiting step of degradation. On the other hand, the relatively slow D1 cleavage might be due to the limited capacity of PSII complexes to migrate to the non-appressed regions of the thylakoids. The repair cycle was proposed to take place in the stroma lamellae or grana margins, to account for the accessibility of damaged D1 to the bulky stromal subunits of FtsH (Andersson and Aro, 2001; Chow and Aro, 2005; Hundal et al., 1990; Uthoff and Baumann, 2018). In such case, the *stt7-9* effect slowing down RC peptides degradation could be due to the impairment in membrane fluidity and movement of damaged PSII RCs.

We have demonstrated that the quenching is lost due to proteolysis (Figs 3, 5), and we propose two mechanisms which can account for that. On one hand, initial cleavage of D1 might result in a loss of q_I_ capacity. The microenvironment of the hypothesized oxidised RC pigment could influence q_I_ or the excitation energy transfer to this site. Another possibility is that the RC complexes with cleaved D1 that still remain assembled (at least a fraction of them is shown in Fig. S10) retain the ability to perform quenching, and only complete degradation of the RC core removes the q_I_ site (see Fig. 7, inner polar plot). However, the quantification of the latter process is difficult due to the complexity of solubilisation of membrane complexes, in particular containing cleaved peptides. If furthermore the RC complex degradation rapidly follows the initial cleavage of D1, distinction between these two processes might prove difficult.

Finally, while we do observe degradation of CP47, this is likely a result of the absence of de novo D1 synthesis (blocked consistently with lincomycin) and thus their assembly partner (de Vitry et al., 1989). Out of RC proteins upon regular photoinhibition, D1 has by far the fastest turnover (Li et al., 2018; Minai et al., 2006; Ohad et al., 1984; Schnettger et al., 1994), confirming that in many cases, replacement of D1 is the only requirement for PSII repair after photodamage.

The experiments in *ftsh1* mutant background provide another advantage in our understanding of photoinhibition-related fluorescence decrease. The stability of observed quenching for at least several hours in the absence of degradation (Figs 3, 5) strongly suggests that q_I_ is the sole quenching mechanism observed, as indicated in the initial timepoints by time-resolved fluorescence (Fig. 1). This is preserved despite the changes in the ultrastructure of thylakoids changes upon photoinhibition (Barbato et al., 1992; Hundal et al., 1990).

### Outlook

In this work, we used integrated approaches to study photoinhibition and the related q_I_ quenching in vivo. A consistent description of all observables across multiple timescales was achieved using a modelling approach in the context of energetic connectivity. We were able to exclude a number of potential quenching sites thanks to spectrally- and time-resolved fluorescence measurements and the use of multiple mutants. We propose that the q_I_ site is formed by a singlet oxygen-mediated attack on one of the PSII RC pigments, simultaneously closing the RC and forming a quenching site. This process at once decreases the F_M_ fluorescence level and raises F_0_. Quenching RCs are then lost through PSII degradation mediated by FtsH. This unified model does not require separate processes to account for the contrasting kinetics of steady-state fluorescence parameters, nor a heterogeneity in quenchers. Our model of photoinhibition where donor- and acceptor-side damage take place independently, with only the latter forming a quenching site, will help understanding this complicated process from a novel perspective.

## Methods

### Strains, culture conditions and photoinhibition treatment

WT strains (CC-1009, CC-1690 (Gallaher et al., 2015)) and *tla3-* (Kirst et al., 2012) were obtained from Chlamydomonas Resource Center. Genetic cross between *stt7-9+* strain (Depège et al., 2003) and *ftsh1-1-* (Malnoë et al., 2014) was performed according to (Harris et al., 1989), yielding the *stt7-9 ftsh1-1* double mutant (Fig. S8). Other strains used (BF4, *pg27* (Bujaldon et al., 2020); *ftsh1-3* (Malnoë et al., 2014); ΔPSII *Fud7.S2+* (Bennoun et al., 1986); ΔPSI *F14.1-* (Dauvillée et al., 2003); were acquired from ChlamyStation (URL: chlamystation.free.fr).

All strains were grown in mixotrophic conditions in shaking glass flasks (Tris-acetate-phosphate medium, 25 °C, continuous 15 μmol photons / m^2^ / s LED light illumination from the bottom, see Fig. S15 for the growth light spectrum). When in early-log phase (1-2 million cells / mL), the cells were spun down at 3000 rpm for 5 min and resuspended in ∼1/5 volume of Tris-phosphate medium (Min) without reduced carbon source. 1 mM NaHCO_3_ was systematically added to replete the bicarbonate binding site in PSII (Brinkert et al., 2016). Concentrated cells were then put back for recovery to their growth chamber for at least 2 h before the measurements.

For photoinhibitory treatment, concentrated cultures were exposed to high light (1500 μmol photons / m^2^ / s; same light source as for growth) in the presence of chloroplast translation inhibitor lincomycin (1.2 mM). The samples from different timepoints after high light treatment were then dark-adapted (15 min) prior to each measurement unless stated otherwise (Figs S1B, S3F). Anoxic photoinhibitory treatment in the same conditions, but starting from at least 30 min before HL exposure the cultures were flushed with a mix of 1% (v/v) CO_2_ in N_2_, and 10 μM DCMU was added prior to photoinhibition. Anoxia after regular photoinhibitory treatment was achieved by an addition of glucose (20 mM) and glucose oxidase (type II from *A. niger*; 50 U / mL) to the cuvette with algae shortly before the measurement.

### Spectroscopy

Fluorescence and transient absorption measurements were done using the JTS-10 apparatus (BioLogic). In fluorescence mode, detecting white LED flashes were filtered through a narrow interference filter (520 nm, 10 nm full width at half-maximum [FWHM], 10 mm) and triggered within short (∼300 μs) dark periods of actinic illumination (‘dark pulse’ mode). The detection was done with a longpass filter (cutoff at 670 nm, 10 mm, Schott). Actinic light was provided from both sides of the cuvette in a custom-build holder, and was set to a subsaturating value of 150 μmol photons / m^2^ / s (630 nm peak) to accurately capture decreasing PSII photochemistry in HL. Saturating pulses of 200 ms provided the same actinic LEDs were used throughout (15 mmol photons / m^2^ / s). The cells were spun during the measurements with a magnetic stirrer. Steady-state fluorescence parameters were corrected to exclude the PSI contribution (3% at F_M_) in data shown in Fig. 5 to account for the absence of PSI in the model membranes.

In absorption mode, P_700_ and plastocyanin redox changes were monitored upon ‘dark pulse’ using detecting flashes passing through a 705 nm (10 nm FWHM) and 730 nm (10 nm FWHM) filters, respectively, and the actinic light was cut from the detectors with Schott RG695 longpass filters. DCMU (10 μM) was added for P_700_ experiments.

PSI:PSII ratio and PSII antenna size were determined as described previously (Nawrocki et al., 2016, 2019; Tian et al., 2019).

Oxygen emission recordings were done simultaneously with fluorescence measurements using a micro-optode (UniSense) stuck inside the culture cuvette. The optode was calibrated in Min medium flushed with air (O_2_ saturated) and in the presence of glucose- and glucose oxidase (anoxia).

Cell absorption spectra were measured using a Cary 4000 spectrophotometer (Varian) fitted with a Diffuse Reflectance Accessory to account for light scattering.

Fluorescence spectra of cells and of BN gel pieces at 77 K were measured with a F-CCDZEN fluorimeter (BeamBio). A LED peaking at 470 nm was used for the excitation.

Time-resolved fluorescence (Fig. 5, S3) was recorded at room temperature with a time-correlated single-photon counting setup (FluoTime 200, PicoQuant). 10 x 10 mm quartz cuvette was used and the cells were stirred using a magnetic stirrer during the measurements. Excitation was provided by a 438 nm laser working at 10 MHz repetition rate and was set to 30 μW using a neutral density filter. Longpass optical filter was placed in front of the detector to prevent scattered light from reaching the photodiode. Detection was done at 680 ± 8 nm with 4 ps bins. DCMU (10 μM) was added to close PSII (F_M_ state). The instrument response function (IRF) was measured through the decay of pinacyanol iodide in methanol (6 ps lifetime; (van Oort et al., 2008)). IRF obtained was 88 ps FWHM in the same conditions as sample measurements. Data analysis was done with the FluoFit software and the decays were deconvoluted with the IRF and 3 exponential decay functions to yield τ_average_ values. In Fig. 5N and 5P, the PSI-related component (∼60 ps) was excluded from the plot to show uniquely the PSII-related contribution to fluorescence decay.

Streak camera measurements (Figs 1, S15-S18) (Van Stokkum et al., 2008) at room temperature were done as described previously (Tian et al., 2019). Sample was stirred during the measurements. 400 nm laser excitation was used (Vitesse Duo, Coherent) and the power at the ∼50 μm diameter spot was set to 15 μW with 250 MHz repetition rate. DCMU (10 μM) was added to close PSII (F_M_ state). Averages of 100 images (10 s integration each) were background- and shading-corrected.

Global analysis of the streak camera images was done as described in (Holzwarth, 1996; van Stokkum, 2018). GloTarAn software was used for the data analysis (Snellenburg et al., 2012), and linked analysis of 3 biological replicas each at 3 timepoints of photoinhibition was performed.

### Biochemistry

Total protein extracts and thylakoids were isolated as described in (Ramundo et al., 2013). Blue native polyacrylamide gel electrophoresis (BN-PAGE) was performed according to (Chen et al., 2019; Wittig et al., 2006). The resolving gel of BN-PAGE used was 4.1-14% T (w/v), while the stacking gel was 3.6% T (w/v), where T stands for the total concentration of acrylamide and bisacrylamide monomers. Blue native gels were cut into strips and loaded onto an SDS (sodium dodecyl sulfonate) gel for the second dimension (Schägger, 2006). 4% T (w/v) was used for the stacking gel and 12% T (w/v) for the resolving gel. After running, the gels were either stained with a Coomassie blue solution (0.1% Coomassie Blue R250 in 10% acetic acid, 40% methanol, and 50% H_2_O) for 2 hr, and de-stained with a de-staining solution (10% acetic acid, 40% methanol, and 50% H_2_O) for 3 x 2 hr; or used for immunoblotting.

Immunoblotting was performed as described in (Nawrocki et al., 2020). 1 μg Chl thylakoid or total protein extract was loaded onto a pre-cast gel (4-12% T (w/v) Bis-Tris Plus, Invitrogen). After electrophoresis, the proteins were transferred to a nitrocellulose membrane. The SDS gel after 2D BN-PAGE was also used for the protein transfer procedure. The membrane was then blocked with 10% (w/v) milk in TTBS (20 mM Tris, pH 7.5, 150mM NaCl, 0.1% Tween 20) for 1 hr and then incubated with different primary antibodies overnight in the dark at 4°C. All antibodies were purchased from Agrisera. The membrane was then washed with TTBS for 4 x 10 min and was incubated with the secondary antibody (goat anti-rabbit IgG) for another 1 hr. The membrane was rewashed with TTBS for 4 x 5 min and was developed for chemiluminescence (with agent SuperSignal West Pico, Thermo Scientific) using a LAS 4000 Image Analyzer (ImageQuant).

### Modelling

The excitation energy transfer rate matrix was constructed using a membrane model originally developed for *A. thaliana* (Amarnath et al., 2016). Excitation-energy transfer was simulated using generalized Forster theory combined with a collection of previously described Hamiltonians and protein structures (Amarnath et al., 2016). Open (k_cs_ = 1.56 ps^-1^, k_rc_ = 0.00625 ps^-1^, k_irr_ = 0.00192 ps^-1^) and closed (k_cs_ = 0.00323 ps^-1^, k_rc_ = 0.00218 ps^-1^) reaction centres had the same electron transport rates as reported previously (Amarnath et al., 2016; Bennett et al., 2013). Damaged reaction centres, which are incapable of charge separation, are either ‘broken’ and have no additional rates of transport, or ‘quenching’ in which case a new loss channel arises with a rate of k_qI_. All calculations presented in the main text use k_qI_ = 0.1 ps^-1^. To construct a rate matrix for a specified number of open, closed, broken, and quenching reaction centres a random number generator was used to determine which of the 128 reaction centres in the membrane belonged to each type. For F_M_ (F_0_) simulations, the number of open (closed) reaction centres is taken to be 0. Finally, the time evolution of the excitation population was simulated using

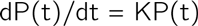

and a fourth order Runge-Kutta integration method with timesteps 0.2 ps. The excitation population was time evolved to 6 ns where residual chlorophyll excitation was negligible. The steady-state fluorescence observables were computed using the simulated fluorescence yield and then averaged over twenty realizations of RC assignments that satisfy the population distribution determined from the kinetic model.

## Acknowledgements

Sandrine Bujaldon is acknowledged for her help with the crossing. This work is supported by a Marie Skłodowska-Curie fellowship xFATE (799083) from the European Commission to W.J.N and by the Netherlands Organization for Scientific Research (NWO; Vici and TOP) to R.C.. X.L. was supported by Chinese Scholarship Council. D.I.G.B. and B.R. acknowledge start-up funds from the Department of Chemistry at Southern Methodist Univeristy. C.d.V. contribution to this work was supported by the Centre National de la Recherche Scientifique and Sorbonne Université (basic support to UMR7141), and by the “Initiative d’Excellence” program (grant “DYNAMO”, Agence Nationale de la Recherche ANR-11-LABX-0011-01).

## Authors contributions

Methodology, W.J.N., B.R., and D.I.G.B.; Software, B.R. and D.I.G.B.; Investigation, W.J.N., X.L., C.H., and C.d.V.; Writing – original draft, W.J.N.; Writing – Review & Editing, all authors; Conceptualization, W.J.N and R.C.; Supervision and Funding acquisition, W.J.N., D.I.G.B., and R.C.

## Supplementary information

**Fig. S1:**
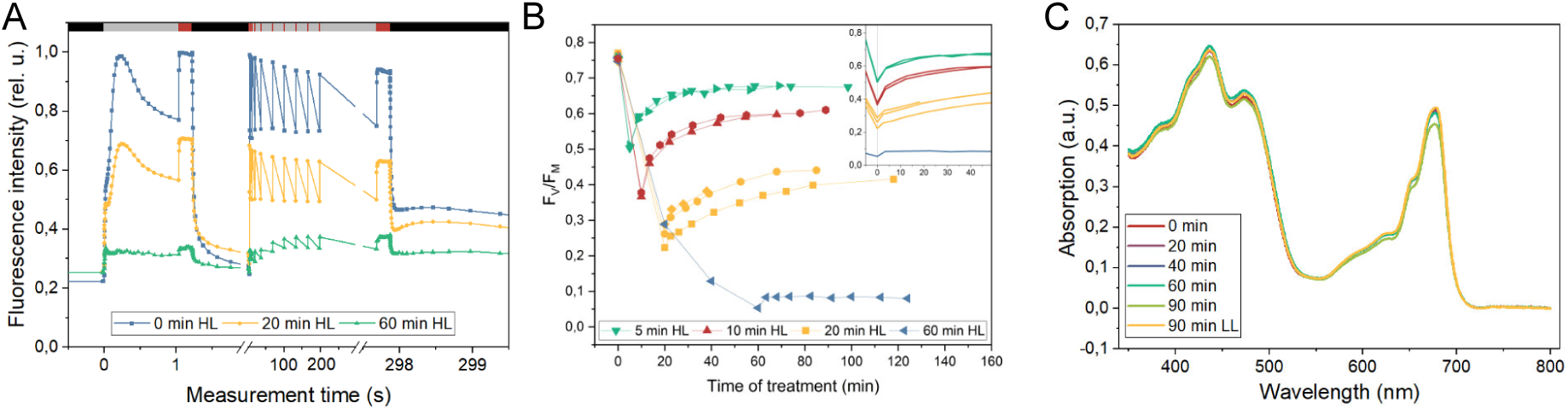
Experimental workflow for q_I_ quenching measurements. All experiments shown were performed in *stt7-9* strain, in the presence of lincomycin which inhibits chloroplast translation and PSII repair. (A) Fluorescence induction protocol used throughout the study and representative raw fluorescence traces at 0, 20, and 90 min HL treatment. Note the non-linear x axis. Black boxes depict dark periods; grey boxes – moderate, non-saturating actinic light illumination (150 μmol photons / m^2^ / s); red boxes – saturating pulses (15 mmol photons / m^2^ / s). (B) Changes in F_V_/F_M_ as a function of HL treatment duration and dark-adaptation following this treatment. Note that about half of the F_V_/F_M_ decrease at short HL timepoints, is rapidly reversible in the dark, indicating that PQ overreduction and not photoinhibition is responsible for this fraction of the variable fluorescence. Inset: the x axis is normalised to 0 upon HL -> dark shift for all traces. Note that ∼15 minutes of dark-adaptation after HL period is sufficient to remove the contribution of the reversible F_V_/F_M_ decrease. (C) Absorption spectra of the cells subjected to photoinhibitory treatment. The spectra were acquired using an integrating sphere. Note that upon photoinhibition, virtually no changes in the absorption were observed, indicating absence of photobleaching in our experimental conditions.

**Fig. S2.**
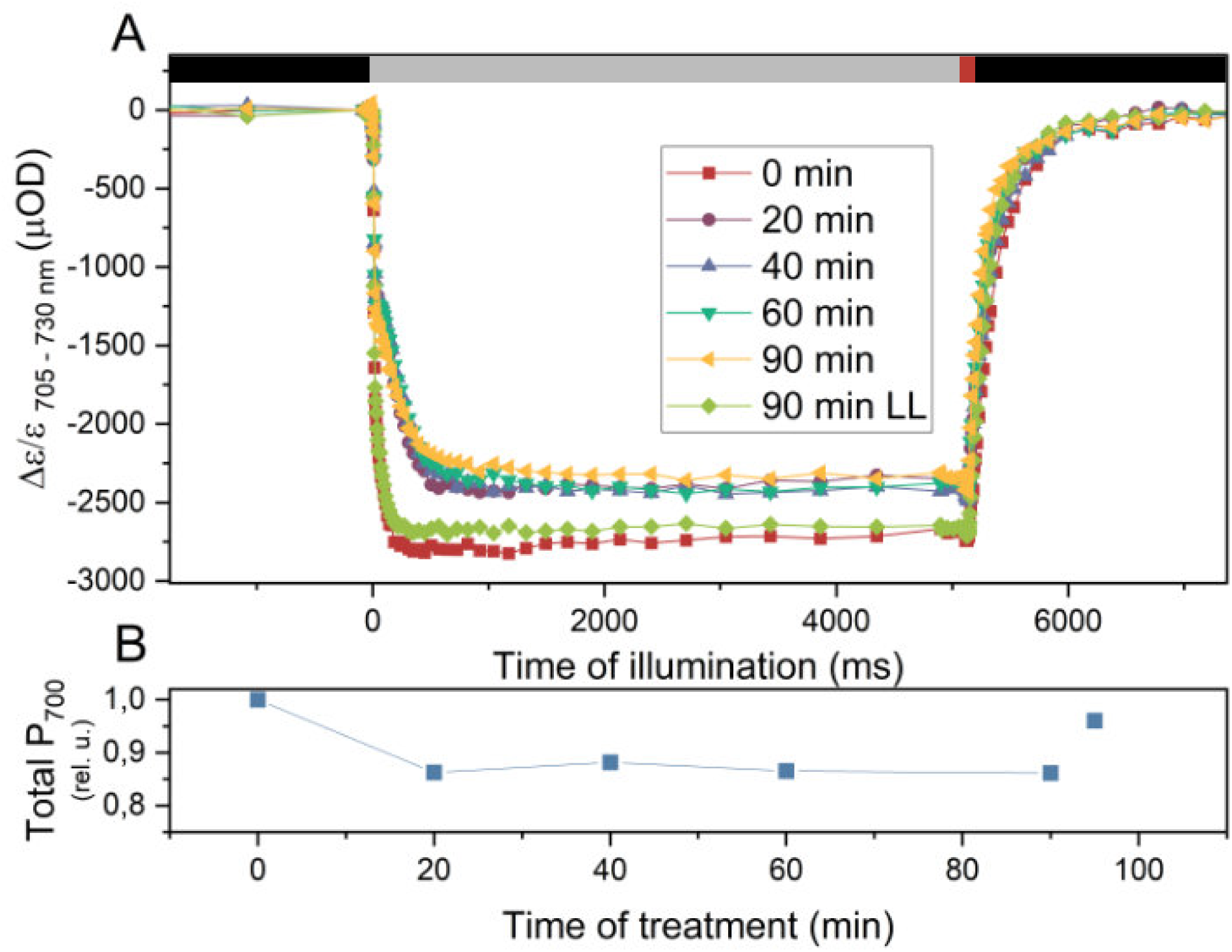
Active P_700_ fraction remains stable throughout the HL treatment. The experiments shown were performed in *stt7-9* strain, in the presence of lincomycin, which inhibits chloroplast translation and PSII repair. (A) Absorption difference plots corresponding to P_700_ oxidation at various timepoints of photoinhibition. The traces were acquired in the presence of DCMU (to obtain the maximal photooxidisable P_700_ quantity), which was added after photoinhibition. Traces obtained at 705 nm were corrected for the plastocyanin signal contribution at 730 nm. Black boxes depict dark period during the protocol; grey boxes – moderate actinic light illumination (150 μmol photons / m^2^ / s); red boxes – saturating pulses (15 mmol photons / m^2^ / s). (B) Quantification of the changes in maximal photoxidisable P_700_ (upon saturating pulse) quantity during photoinhibition. The timepoint at 95’ is a control not exposed to HL.

**Fig. S3.**
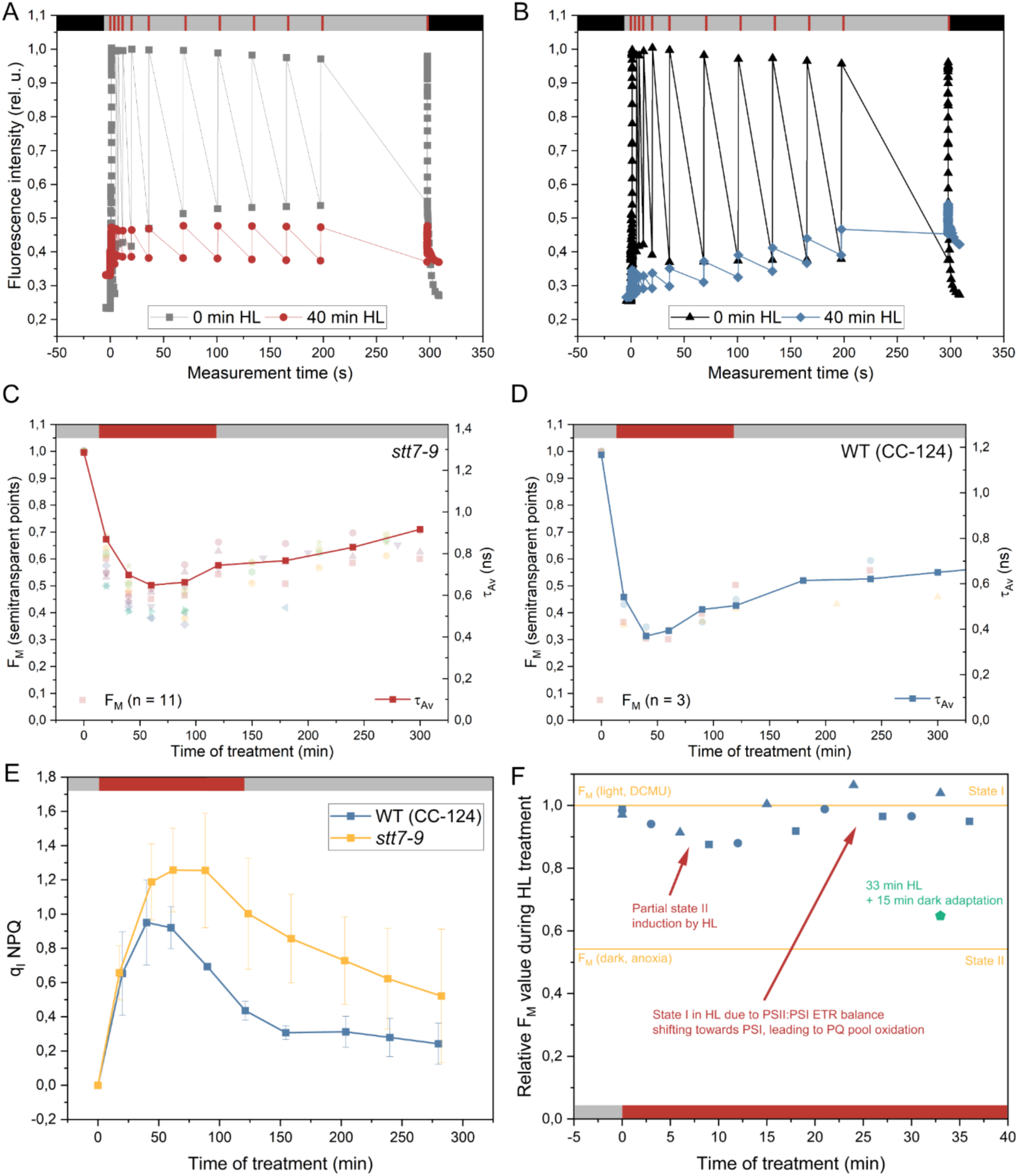
Disentangling q_I_ quenching and State transitions from steady-state and time-resolved fluorescence measurements. All experiments shown were performed in the presence of lincomycin which inhibits chloroplast translation and PSII repair. (A) Fluorescence induction traces at 0- and 40 min HL treatment in the *stt7-9* strain. Note the unchanged level of F_M_ (red boxes) throughout the illumination. Black boxes depict dark period during the protocol; grey boxes – moderate, non-saturating actinic light illumination (150 μmol photons / m^2^ / s); red boxes – saturating pulses (15 mmol photons / m^2^ / s). (B) Like (A) but in the WT (CC-124) strain. Note the stable behaviour of F_M_ during illumination after 0 min HL, and the slow rise of F_M_ during the light period after the algae were exposed to 40 min HL and dark-adapted for 15 minutes. This rise corresponds to a State II -> State I transition. (C) F_M_ values evolution upon HL treatment in *stt7-9* strain. The τ_Av_ value is the amplitude-weighted average lifetime at F_M_ (in the presence of DCMU) obtained using TCSPC (broadband detection at 680 ± 8 nm, see Materials and methods for details). Note the excellent correlation between the F_M_ and τ_Av_ throughout the treatment, indicating that in experimental conditions where photobleaching or State transitions do not take place, steady-state fluorescence can be used as an approximation of q_I_ quenching. Grey boxes – low light illumination (15 μmol photons / m^2^ / s); red boxes – HL period (1500 μmol photons / m^2^ / s). (D) The same as (C) but in WT (CC-124) strain. Note that the fluorescence values at early photoinhibition timepoints are lower in the WT background compared to *stt7-9*. This behaviour is due to the quenching of LHCII in State II. The last F_M’_ value is therefore defined as F_M’(State I)_ as during the illumination, a transition from STII to STI took place. (E) Influence of STT7 on quenching kinetics and loss of q_I_. In experiments which include the low-light fluorescence induction (such as in (A)), it is possible to obtain F_M’(State I)_ value in the WT background. The plot shows the q_I_ behaviour (calculated as [(F_M_-F_M’(State I)_)/ F_M’(State I)_]) in the WT (CC-124) and *stt7-9* strains. Note that the initial kinetics of photoinhibition are identical in the two strains, but that the loss of q_I_ is faster in WT background. Grey boxes – low light illumination (15 μmol photons / m^2^ / s); red boxes – HL period (1500 μmol photons / m^2^ / s). (F) WT strains are virtually fully in State I under high light treatment. The plot shows a photoinhibition experiment with no dark-adaptation time between the HL treatment and fluorescence measurement. Shown is the quantification of amplitude between the F_M(initial)_ (corresponding to 0 s in (B)) and F_M’(State I)_ (300 s in (B)) for early HL treatment timepoints (blue). The maximal- and minimal F_M_ values (yellow lines) were determined by an illumination with DCMU (complete State I) and by anoxic treatment in darkness (full State II). Note that the F_M(initial)_, and thus the state of WT cell in high light, is almost completely at the level corresponding to State I. The green timepoint at 33 min includes dark-adaptation and substantiates the interpretation that cells following HL treatment shift to State II upon dark-adaptation (compare with panel (B)). Grey boxes – low light illumination (15 μmol photons / m^2^ / s); red boxes – HL period (1500 μmol photons / m^2^ / s).

**Fig. S4.**
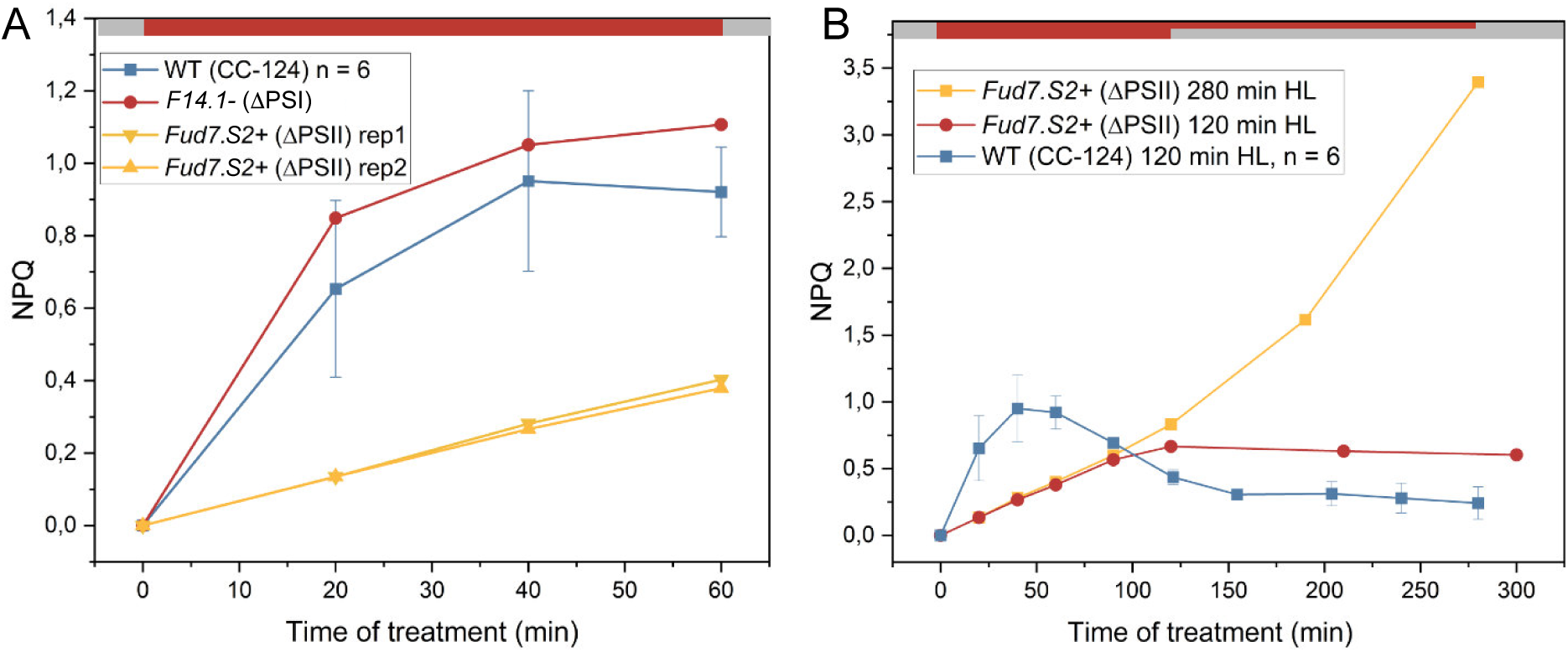
HL-induced quenching in PSII- and PSI RC mutants. All experiments shown were performed in the presence of lincomycin which inhibits chloroplast translation. Grey boxes – low light illumination (15 μmol photons / m^2^ / s); red boxes – HL period (1500 μmol photons / m^2^ / s). (A) NPQ values in reaction centres mutants and the reference WT (CC-124) strain during the HL treatment. q_I_ NPQ was calculated as [(F_M_-F_M’(State I)_)/ F_M’(State I)_] (see Fig. S3). (B) NPQ values in PSII-lacking strain at longer illumination times. Notice that the decrease of fluorescence (apparent increase of NPQ) is continuous upon HL treatment in *Fud7.S2+*, contrary to the situation in WT (Fig. S6)

**Fig. S5.**
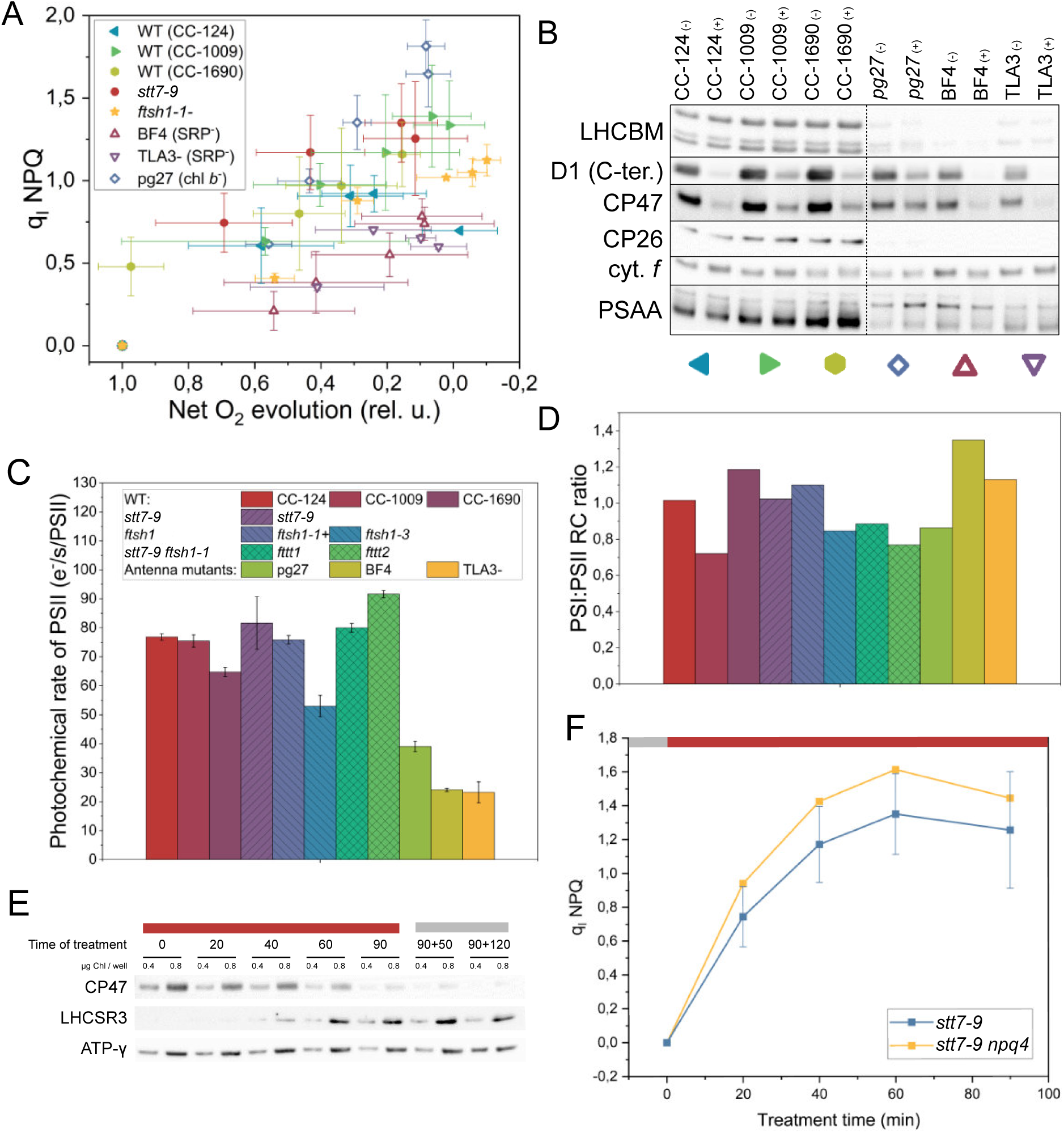
q_I_ does not take place in the antennae. All experiments shown were performed in the presence of lincomycin. (A) q_I_ in antenna mutants and other reference strains. To account for the variable rates of the acceptor-side photoinhibition in the PSII antenna mutants (due to lower effective photochemical rate of PSII during illumination), the development of the quenching is shown as a function of decreasing O_2_ evolution capacity rather than as a function of time. (B) Immunoblot of selected thylakoid membrane proteins in WT strains and in light harvesting mutants before (-) and after (+) 90 min HL treatment in the presence of lincomycin. The coloured symbols correspond to those in panel (C). Note that Agrisera **AS09 408** ⍺-LHCBM antibody used recognizes all major LHCII in Chlamydomonas. (C) “Antenna size” (maximal light-limited photochemical rate) of PSII (D) PSI:PSII reaction centre ratio measured in vivo. (E) Accumulation of LHCSR3 during photoinhibition in the *stt7-9* strain. CP47 is shown to highlight the ongoing photoinhibition, and the gamma subunit of ATP synthase is shown as a loading control. Grey boxes – low light illumination (15 μmol photons / m^2^ / s); red boxes – HL period (1500 μmol photons / m^2^ / s). (F) q_I_ induction in *stt7-9 npq4* strain lacking LHCSR3. Grey boxes – low light illumination (15 μmol photons / m^2^ / s); red boxes – HL period (1500 μmol photons / m^2^ / s).

**Fig. S6.**
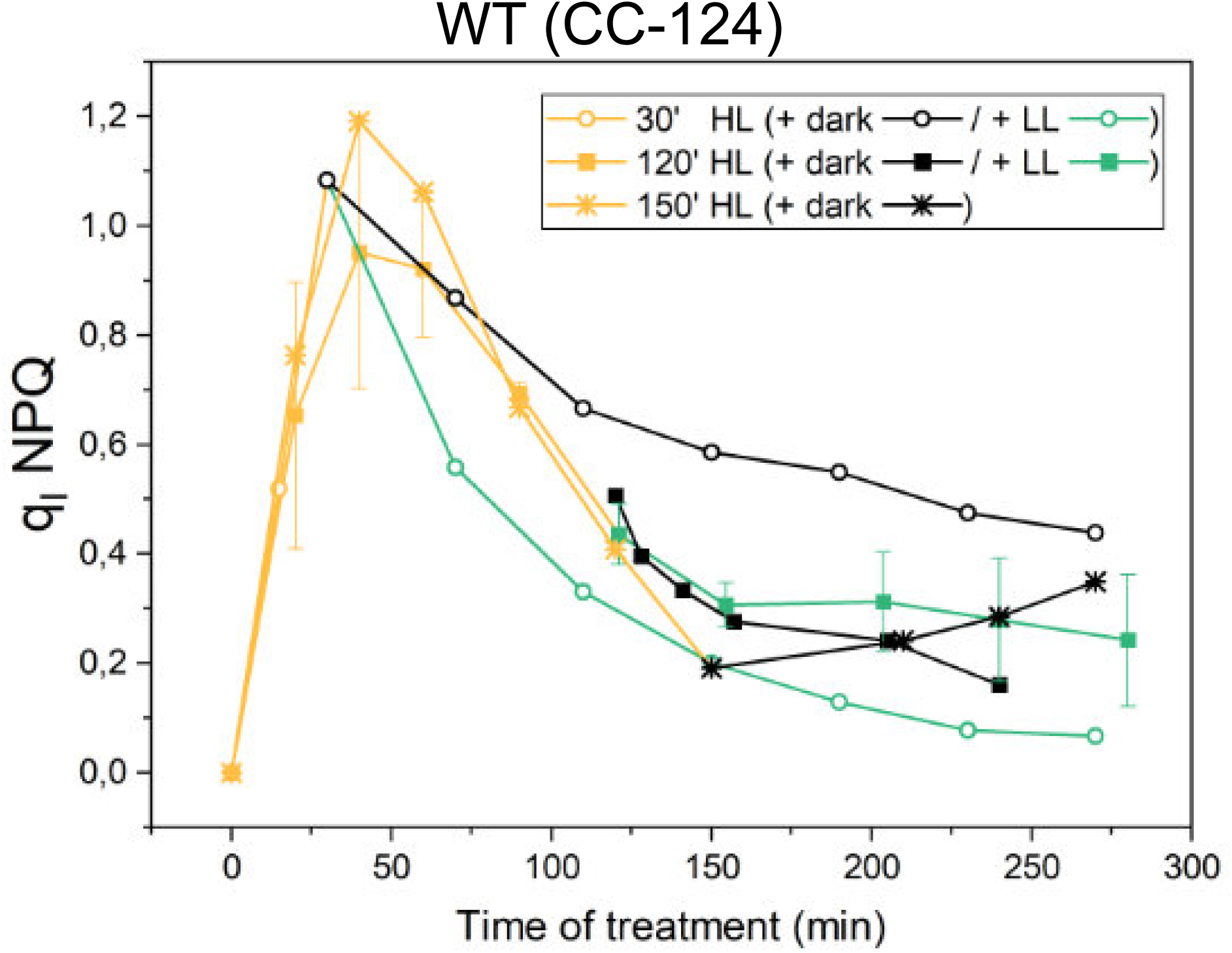
Loss of q_I_ is independent of light. Shown experiments were performed in the presence of lincomycin. q_I_ was monitored upon WT (CC-124) exposure to three different durations of HL and upon ‘recovery’ in low light or darkness. Note that the decrease of quenching is observed in all conditions.

**Fig. S7.**
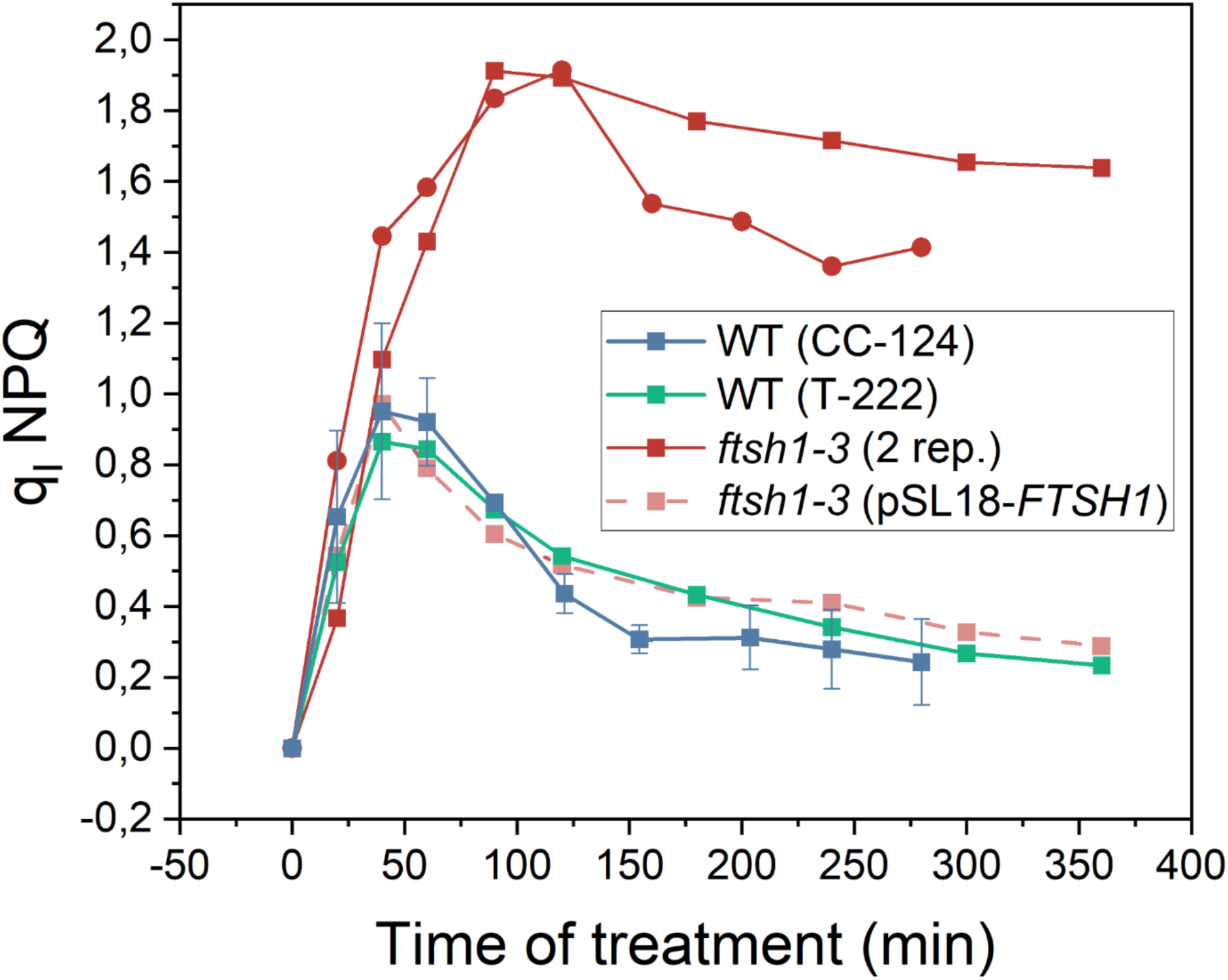
q_I_ in *ftsh1-3*. Shown experiments were performed in the presence of lincomycin. *ftsh1-3*(pSL18-*FTSH1*) is the *ftsh1-3* strain complemented with the genomic sequence of *ftsh1*.

**Fig. S8.**
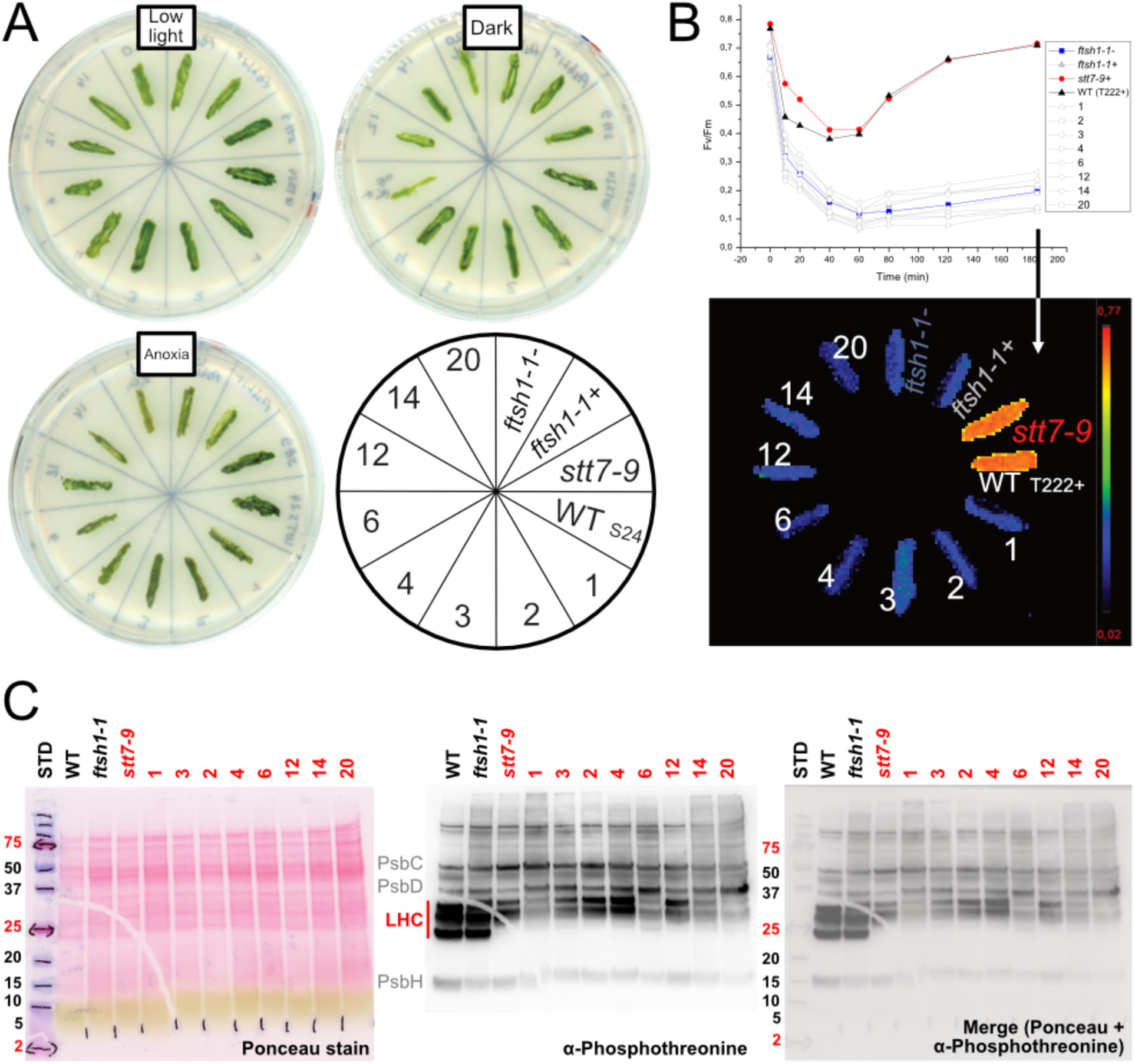
Construction and selection of the double *stt7-9 ftsh1-1* mutants. (A) Macroscopic phenotype of the cross descendants from *ftsh1-1* and *stt7-9* grown on solid media in different conditions. (B) Recovery from photoinhibition is impaired in the double *stt7-9 ftsh1-1* mutants. Plot of F_V_/F_M_ upon 60-min photoinhibition and 120 min recovery in low light, in the absence of lincomycin, on plates. Below the F_V_/F_M_ image in each colony used for the quantification. (C) *stt7-9 ftsh1-1* mutants are impaired in state transitions. Whole cell extracts isolated from the cross descendants were subjected to an SDS-PAGE and immunoblotting with an α-phosphothreonine antibody. Note that the LHCII bands are not phosphorylated in the descendants.

**Fig. S9.**
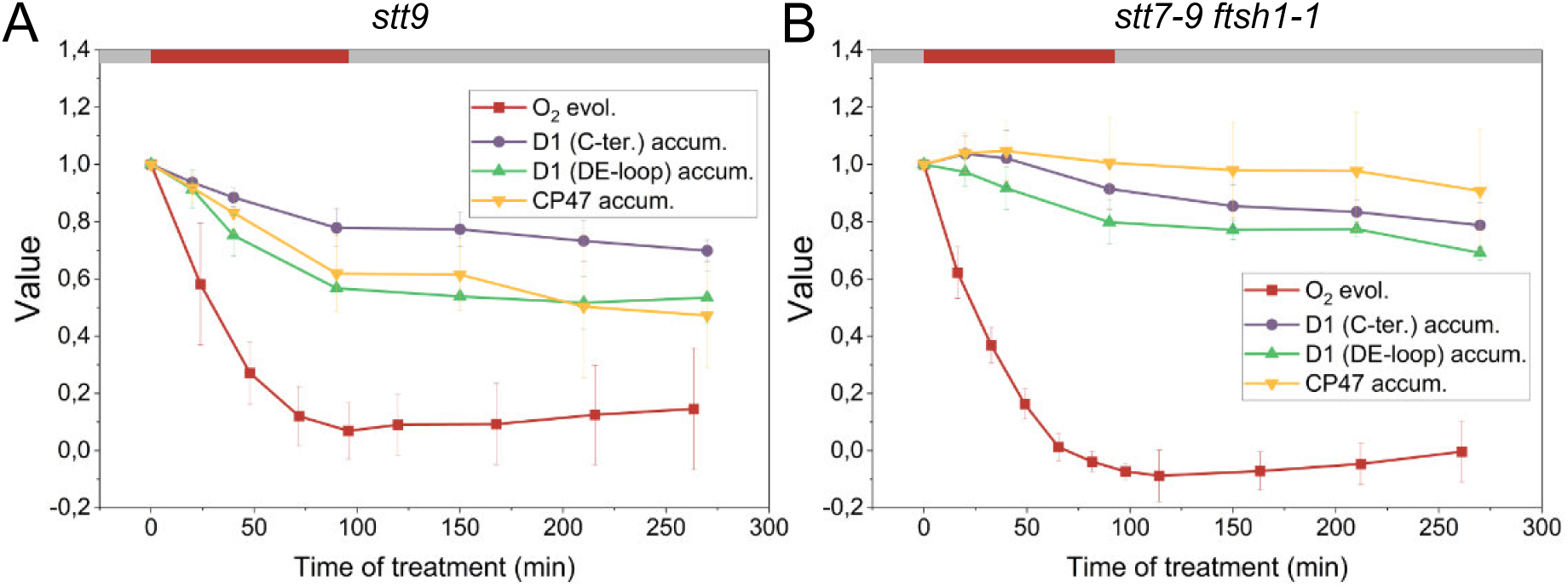
Quantification of full-length PSII RC proteins loss upon photoinhibition. The experiments shown were performed in the presence of lincomycin. The immunoblot (Fig. 3) signal of each protein was normalized to its signal in the 0 min HL sample. Grey boxes – low light illumination (15 μmol photons / m^2^ / s); red boxes – HL period (1500 μmol photons / m^2^ / s). Means ± SD are shown from n = 3 biological replicas (protein accumulation) and n = 6 (oxygen evolution capacity). (A) *stt7-9* (B) *stt7-9 ftsh1-1*

**Fig. S10:**
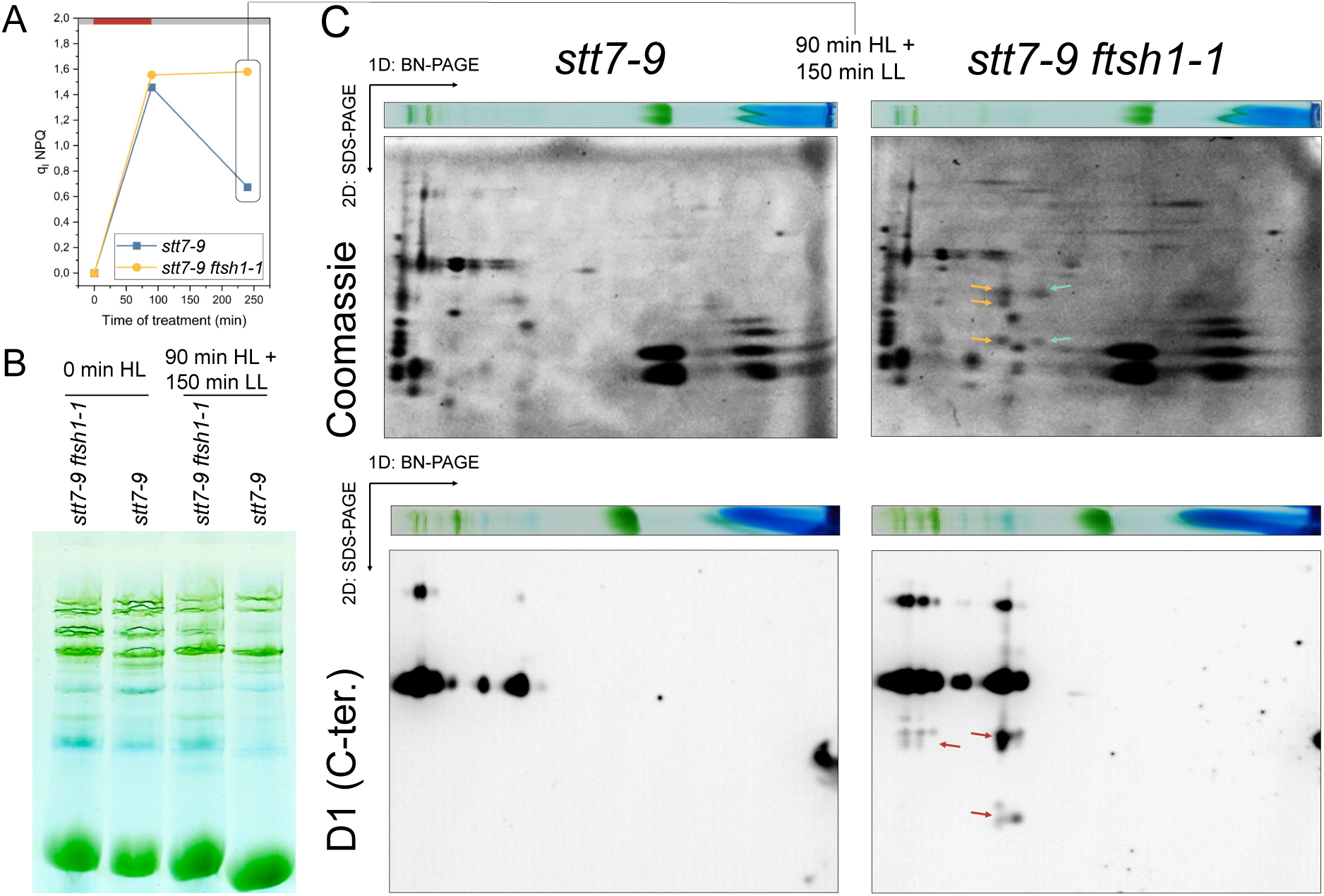
PSII supercomplexes with cleaved D1 accumulate in the *ftsh1* background. All experiments shown were performed in the presence of lincomycin. (A) q_I_ kinetics in the samples used for BN-PAGE and 1D-BN-PAGE/2D-SDS-PAGE shown in the panels (B) and (C). Red boxes – HL illumination (1500 μmol photons / m^2^ / s); grey boxes – low light period (15 μmol photons / m^2^ / s). (B) BN-PAGE of *stt7-9* and *stt7-9 ftsh1-1* strains prior to- and following HL treatment and recovery from q_I_. (C) 1D-BN-PAGE/2D-SDS-PAGE of the samples from (B). Top, Coomassie staining of the gels. Bottom, α-D1 (C-ter.) immunoblotting. Arrows indicate regions of interest: CP47, CP43, and D2 bands (orange), CP43 and D2 bands (green); and fragments of D1 (red). Note that the D1 fragments present in the *stt7-9 ftsh1-1* mutant remained to a certain extent a part of PSII complexes.

**Fig. S11.**
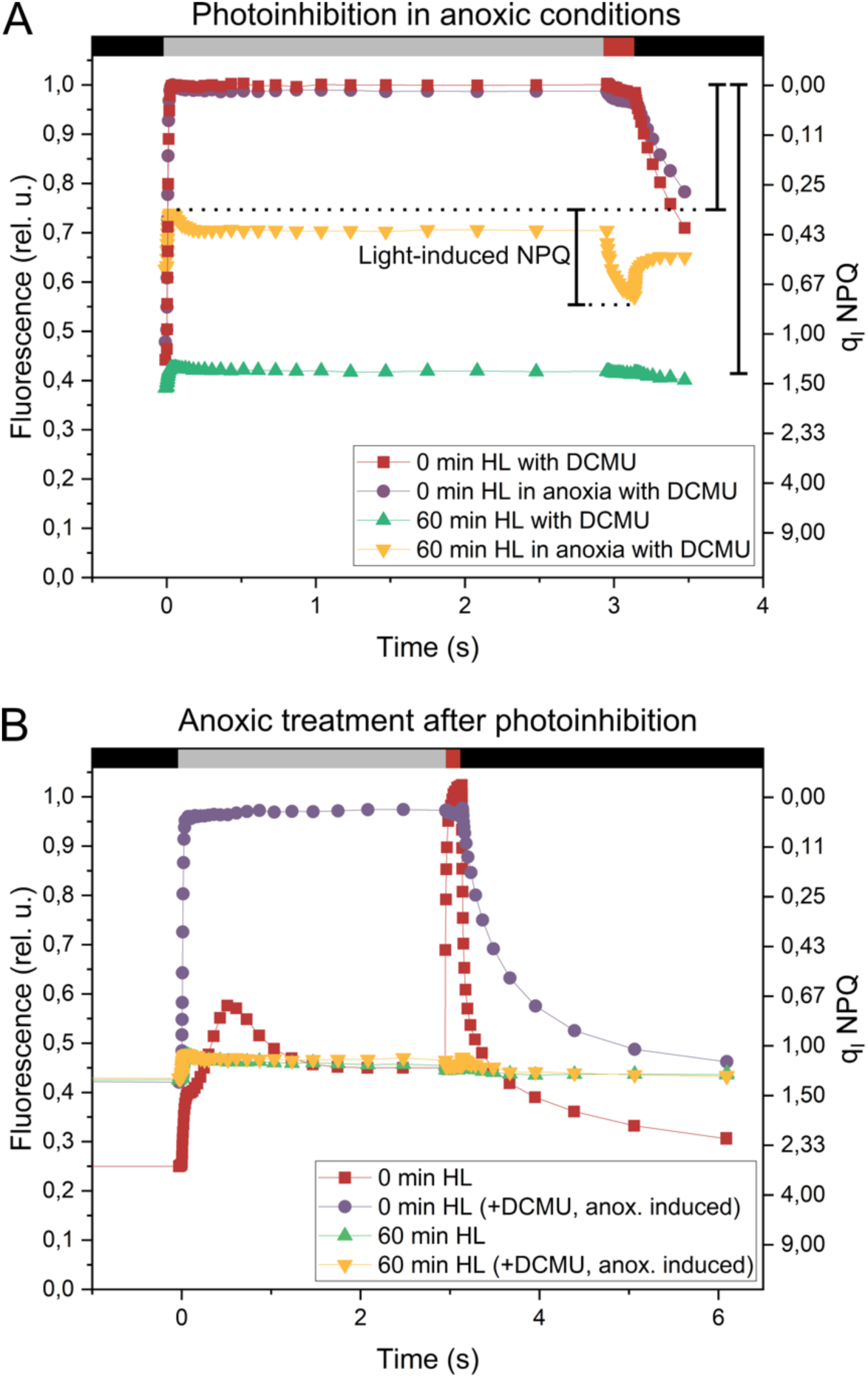
Oxygen-dependence of q_I_ formation and oxygen-dependence of q_I_ quenching. All experiments shown were performed in the *stt7-9* strain, in the presence of lincomycin. Black boxes depict dark period during the protocol; grey boxes – moderate actinic light illumination (150 μmol photons / m^2^ / s); red boxes – saturating pulses (15 mmol photons / m^2^ / s). (A) Representative raw fluorescence traces of cells subjected to anoxic- and oxic photoinhibition in the presence of DCMU. Note the appearance of light-induced quenching in the cells where photoinhibition was induced in anoxia. (B) Fluorescence kinetics of cells where anoxia was induced after photoinhibitory treatment. Note that presence of oxygen in photoinhibited cells has no effect on the amplitude of fluorescence.

**Fig. S12.**
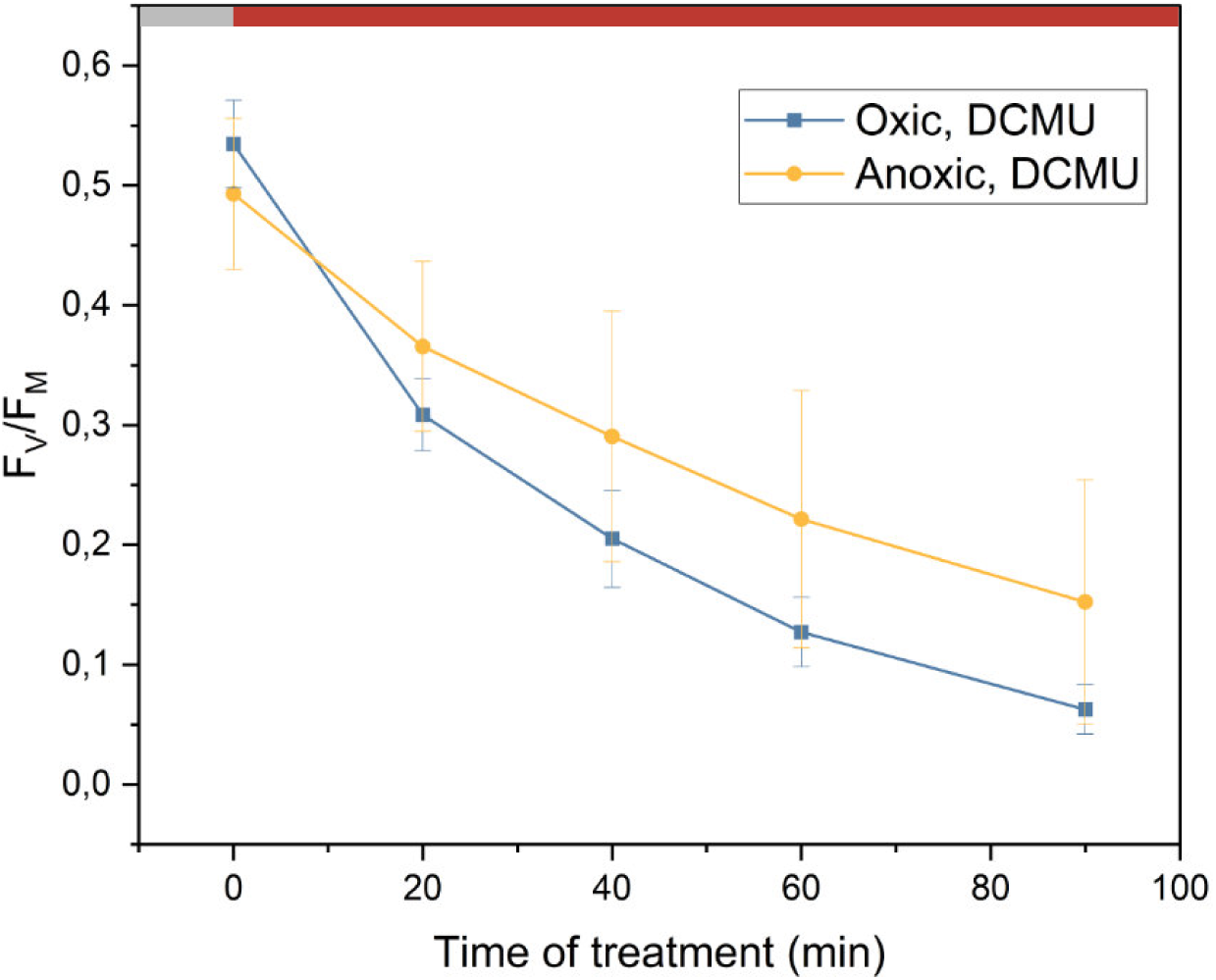
F_V_/F_M_ upon photoinhibition in anoxia. The experiments were performed on the *stt7-9* strain, in the presence of lincomycin. Grey box – low light illumination (15 μmol photons / m^2^ / s); red boxes – HL period (1500 μmol photons / m^2^ / s).

**Fig. S13.**
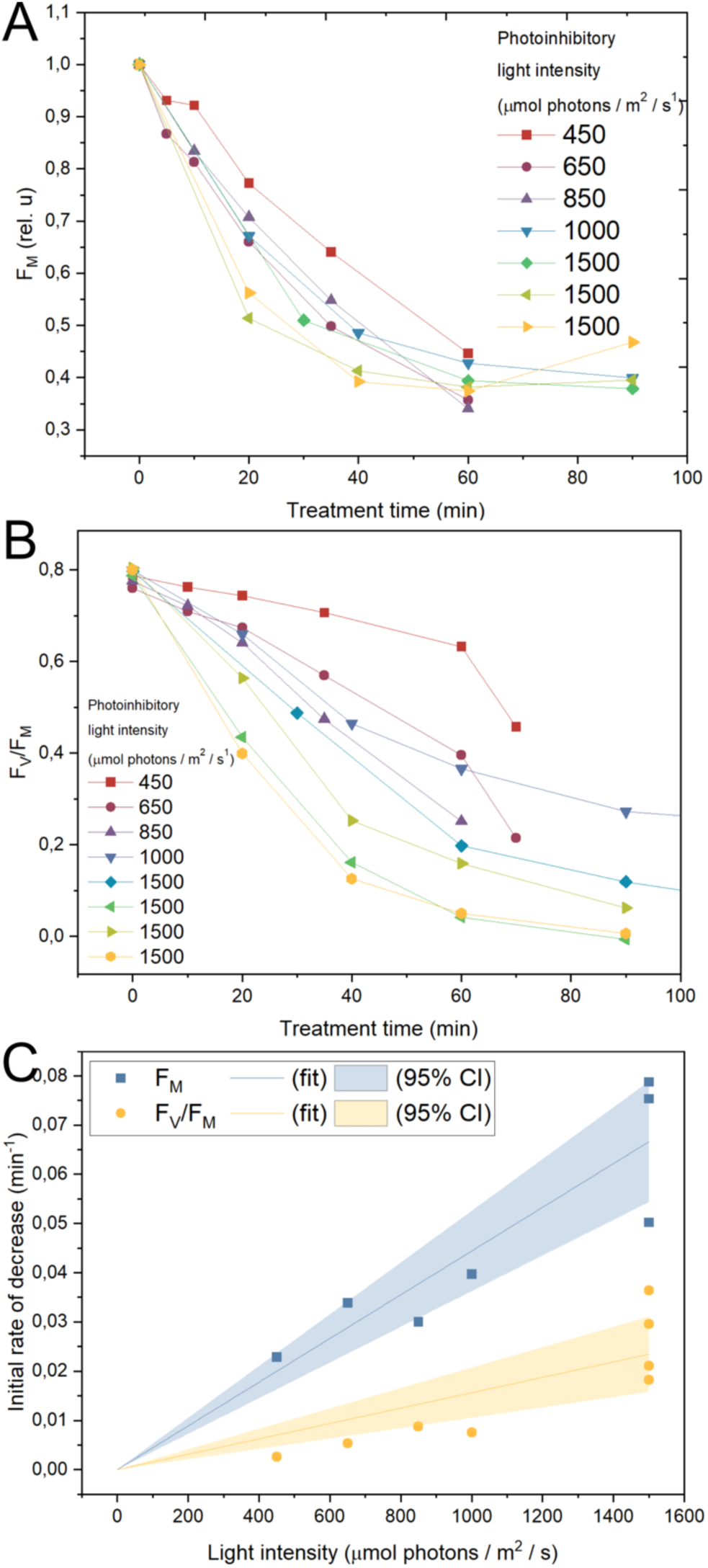
Corresponding kinetics of F_M_ and F_V_/F_M_ upon photoinhibition at various light intensities. All experiments shown were performed in the *stt7-9* strain, in the presence of lincomycin which inhibits chloroplast translation. Selected traces shown for clarity. (A) F_M_; (B) F_V_/F_M_; (C) Quantification of the initial rate of changes from (A) and (B). Shaded areas represent the 95% confidence intervals from linear fits of the data forced through the intercept.

**Fig. S14.**
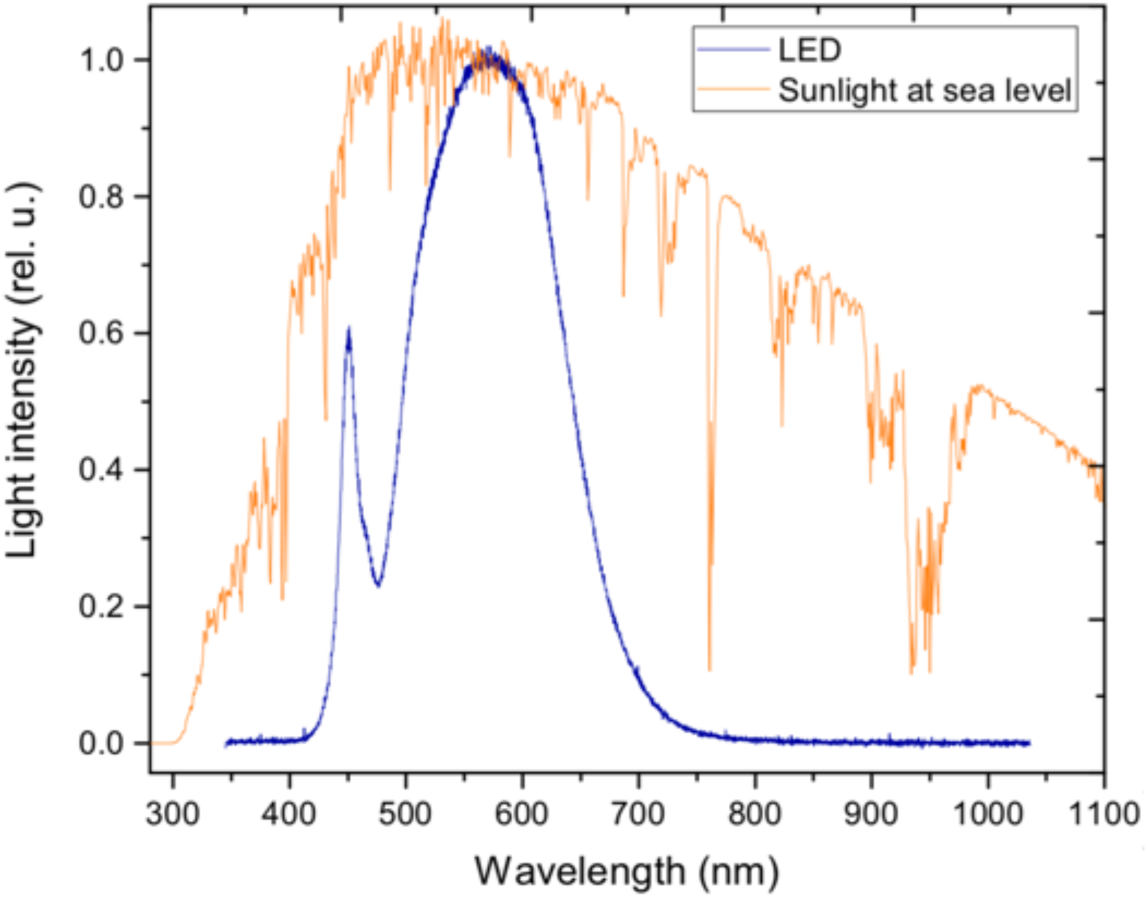
Light spectrum of the sun and of the LED used to grow Chlamydomonas and for the photoinhibitory treatment.

**Fig. S15.**
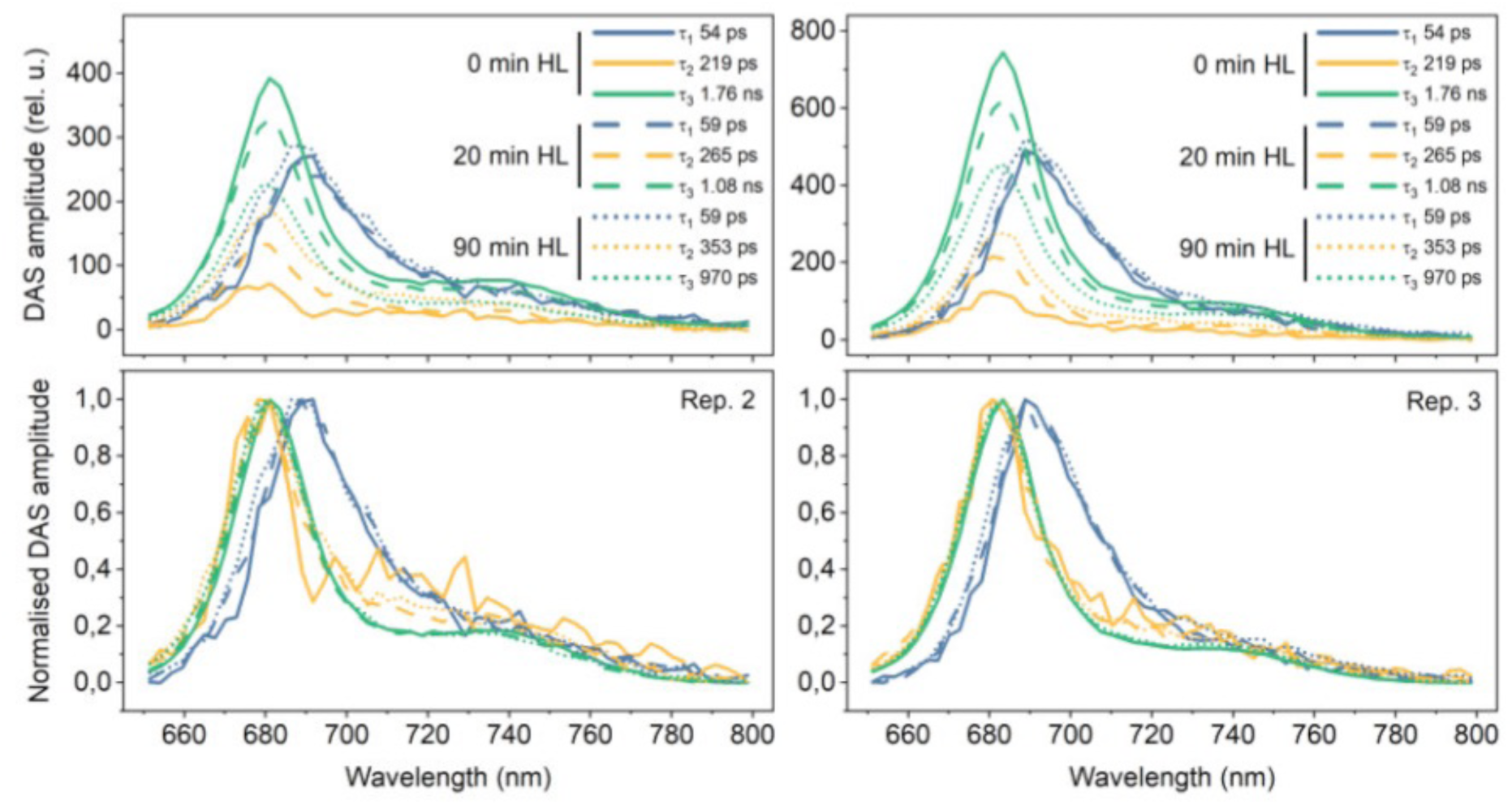
Decay associated spectra of 2 independent repeats of time-resolved measurements of *stt7-9* strain during photoinhibition performed with a streak camera setup. The experiments were done in the presence of lincomycin which inhibits chloroplast translation. The marginal increase in PSI-related component lifetime and amplitude was earlier proposed to stem from a residual capacity for State transitions in the *stt7-9* mutant (Bergner et al., 2015; Tian et al., 2019).

**Fig. S16.**
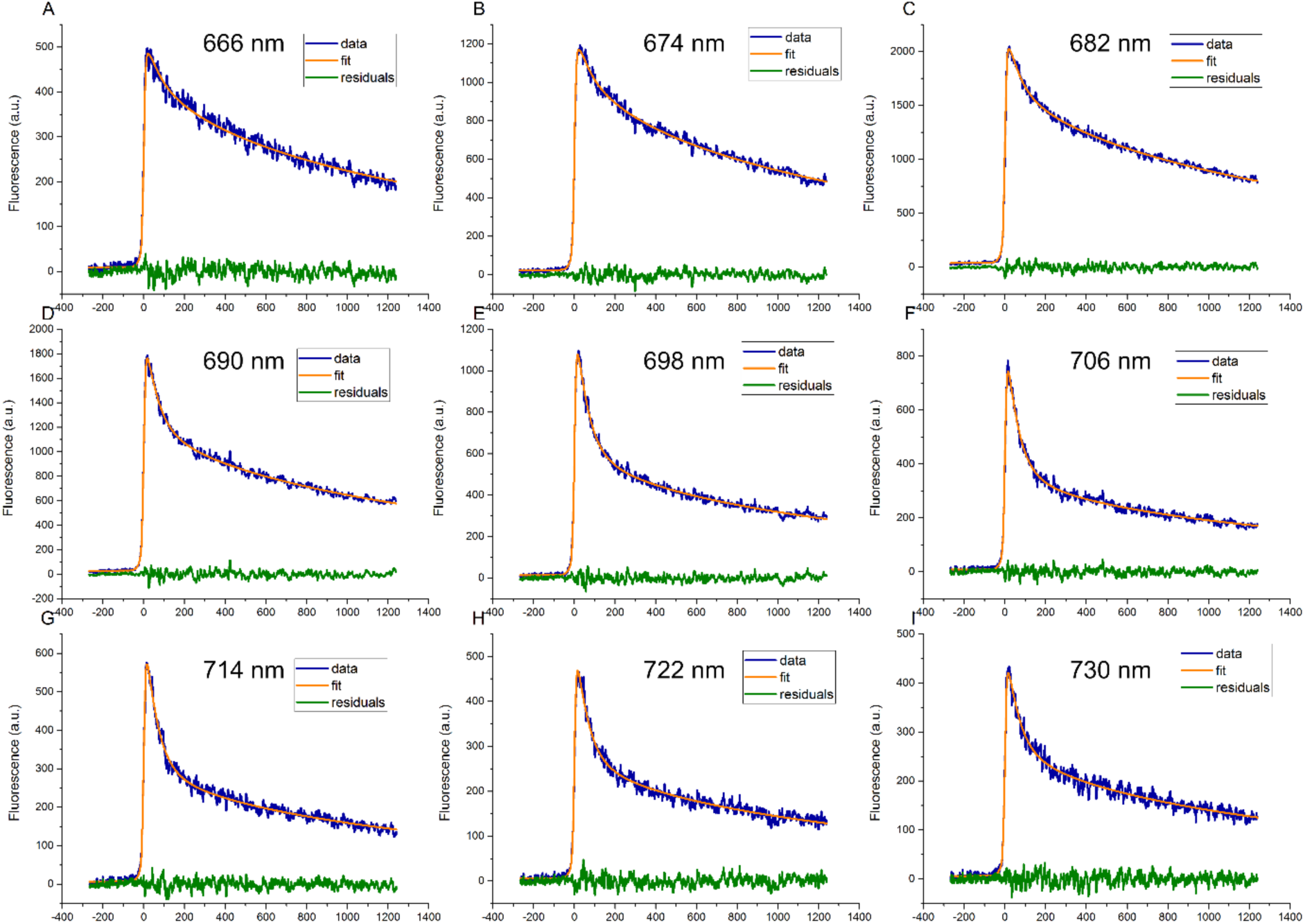
Time-resolved fluorescence traces, the fitted curves, and their residuals at selected wavelengths. Data for 0 min HL treatment shown.

**Fig. S17.**
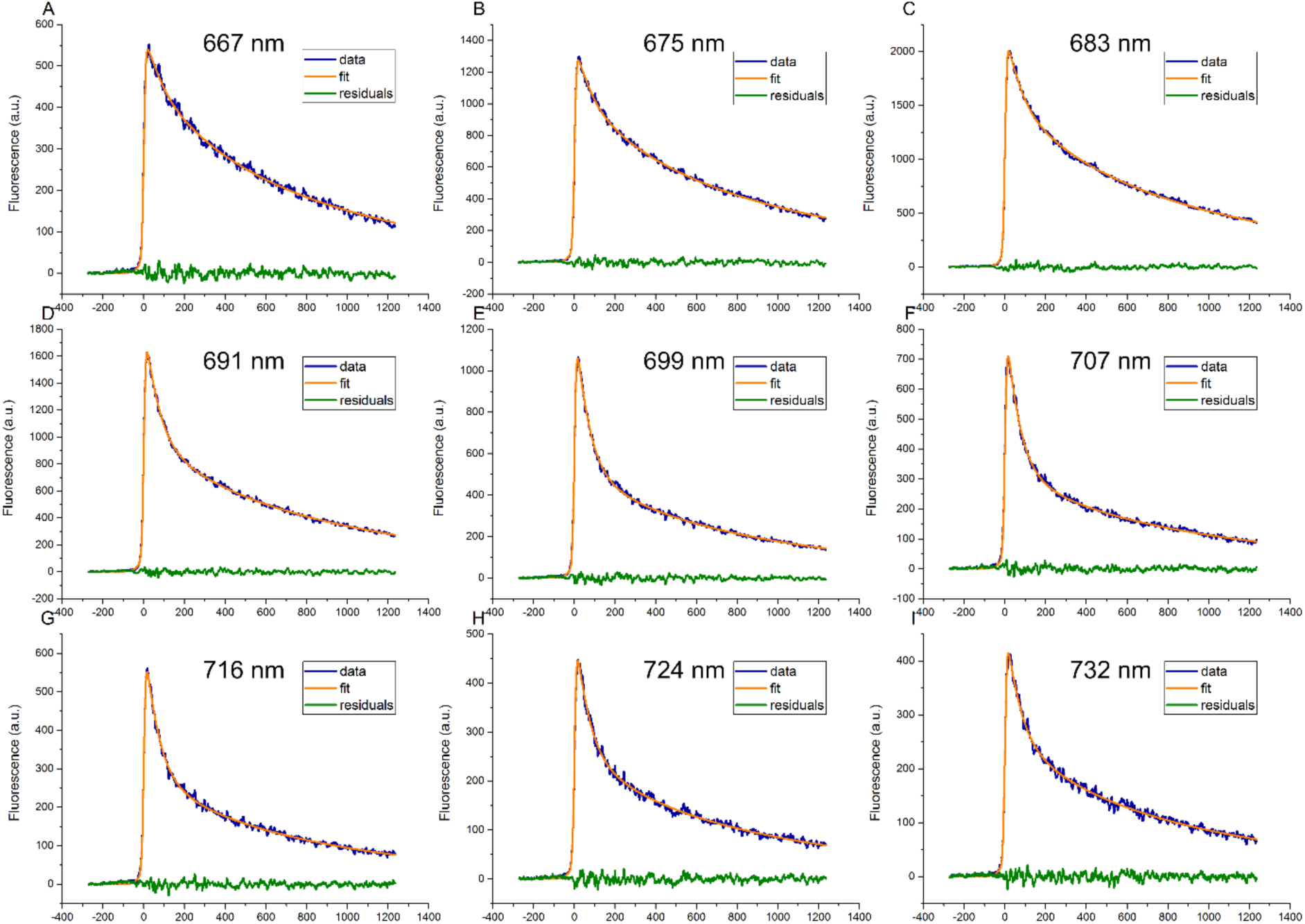
Time-resolved fluorescence traces, the fitted curves, and their residuals at selected wavelengths. Data for 20 min HL treatment shown.

**Fig. S18.**
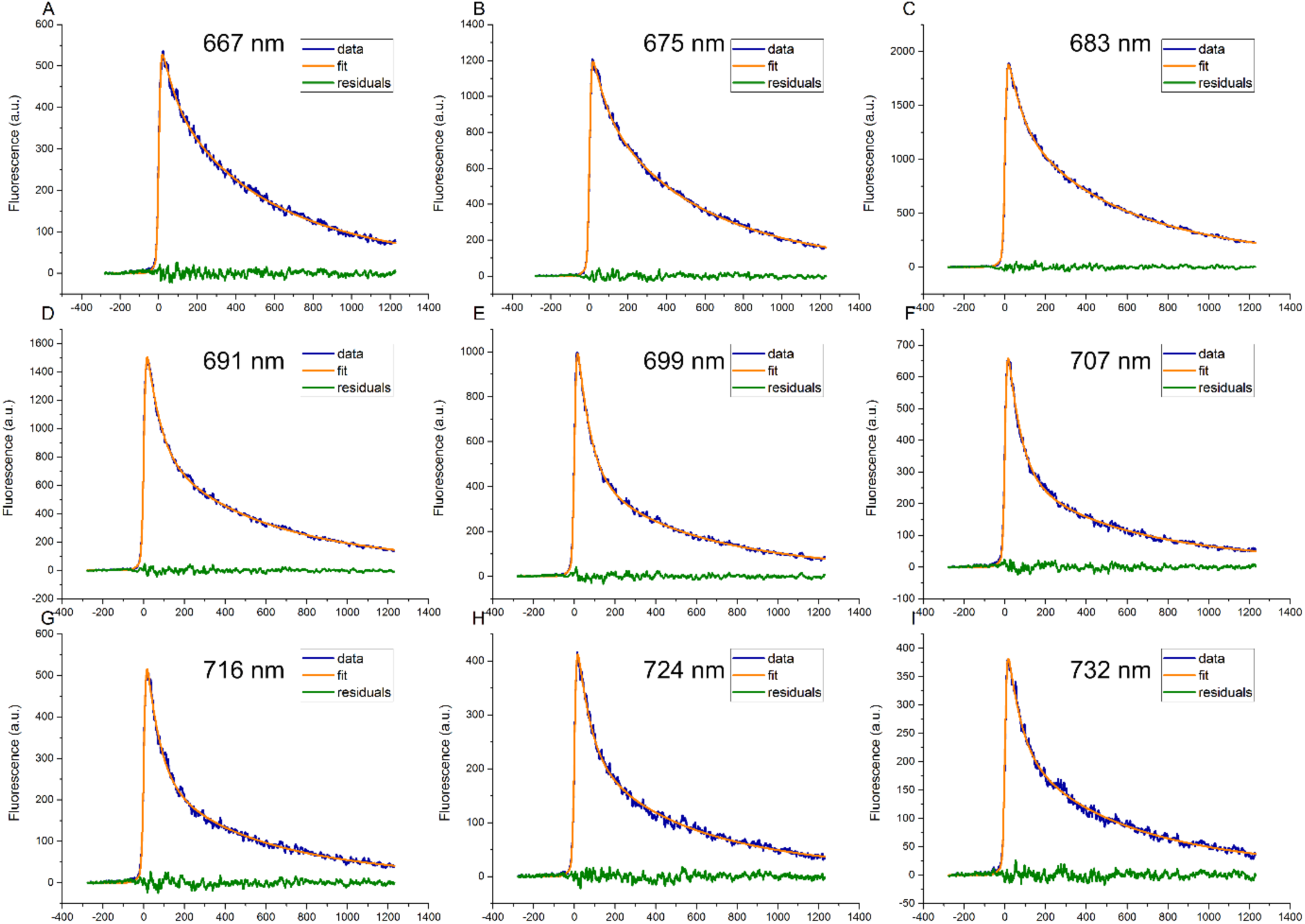
Time-resolved fluorescence traces, the fitted curves, and their residuals at selected wavelengths. Data for 90 min HL treatment shown.

### Supplementary discussion

#### Fitting of kinetic parameters

The parameters defining the kinetic model (Fig. 5), i.e. the kinetic rates between varying RC types, were fit to reproduce the steady state fluorescence measurements (F_0_, F_M_, and F_V_/F_M_). For each set of kinetic parameters, the squared error

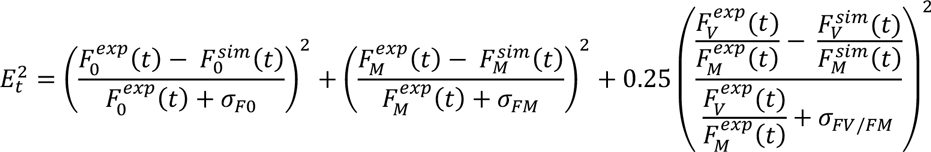

was determined at every timepoint for both the *stt7-9* mutant and the *stt7-9 ftsh1-1* double mutant strains. We used the root-mean squared error across the two strains to describe the quality of fit. We included the F_V_/F_M_ term in order to bias the fit towards solutions that reproduced the relative F_0_ and F_M_ behaviour. We also added the average standard deviation of each experimental measurement (*σ_X_*) to the denominator of the error in order to decrease the sensitivity of the fit in proportion to the uncertainty in the measurement.

The four-parameter kinetic models (described below) showed a rugged error landscape. As a result, we determined our optimal parameters using the dual annealing minimizer with a gradient descent optimization method implemented in the Scipy optimize library (Virtanen et al., 2020).

#### Testing Different Kinetic Models

The linear model presented in Fig. 5 succeeds in fitting the experimental data. Other, more elaborate kinetic models will also reproduce this data. We have tested different kinetic models with four or fewer parameters and found they provide the same or lower quality reproduction of the experimental data. Fig. S19A (left) shows the three-parameter kinetic model used for calculations presented in the main text, denoted as the linear model. In this model, the rate between quenching and broken RCs was allowed to vary between the two strains which we depict schematically as two arrows between those boxes in the kinetic diagram. We compared the three-parameter model to additional models of higher complexity by calculating the minimal error for each kinetic model across a range of quenching rates (Fig. S19B). In particular, we hypothesized that there could be a light induced mechanism that directly converts active to broken RCs (Fig. S19A, right). We then considered a four-parameter model which, like the linear model, allowed the rate from quenching to broken RCs to vary between the two strains (Model-4A). The linear model shows a sharply defined minimal error for a quenching rate (k_qI_) of 0.1 ps^-1^, while model-4A has essentially equivalent error to the linear model for low quenching rates (k_qI_ <0.1 ps^-1^) but returns similarly small errors for a broad range of quenching rates. The lack of any substantial improvement in the overall fit to the fluorescence measurements in Model-4A is notable given the greater flexibility available for fitting. We also tested other four-parameter kinetic models where the rates from active to quenching RCs and active to broken RCs were allowed to change between the two mutant strains (Model-4B/C, Fig. S19A right). Strikingly, the overall error for both of these models increased compared to the simpler linear model. This suggests that the flexibility in describing the connection between quenching and broken reaction centres is critical to describing the experimental observations.

**Fig. S19.**
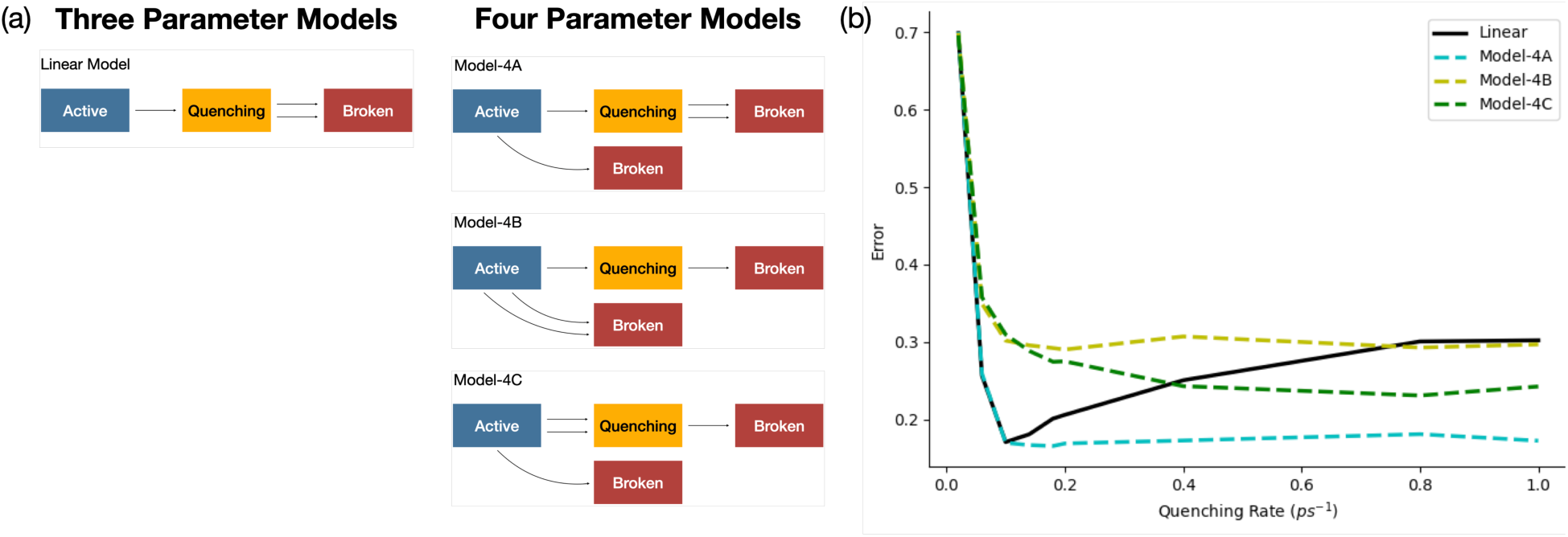
Kinetic Models and Error Analysis. (A) Schematic diagram of the four different kinetic models tested. (B) Overall error of each kinetic model as a function of the quenching rate (k_qI_).

